# Stage-Dependent Requirement for SETX in Human Motor Neuron Maturation and Survival

**DOI:** 10.64898/2026.11.17.767329

**Authors:** Md Abir Hosain, Md Kobirul Islam, Yuntian Duan, Haochen Cui, Xinggui Shen, Tengyu (Danny) Ko, Shisheng Li, Baojin Ding

## Abstract

Senataxin (SETX) is an RNA/DNA helicase involved in transcription, RNA processing, R-loop resolution, and genome maintenance. Mutations in *SETX* cause neurological disorders, but its role in human motor neuron (MN) development remains unclear. Here, we examined SETX function using isogenic SETX knockout (KO) human induced pluripotent stem cells (hiPSCs), shRNA-mediated knockdown, and direct conversion of fibroblasts into MNs. SETX expression increased during differentiation from hiPSCs to neural progenitor cells (NPCs) and MNs. SETX loss slowed hiPSC and NPC proliferation without preventing MN generation. However, SETX-deficient MNs developed shorter neurites, dystrophic swellings, and progressive neuronal loss. MEA recordings showed reduced firing and burst activity and almost no synchronized network bursts. RNA hybrid accumulation increased after SETX loss, with the largest relative increase in MNs among the cell types examined. RNA-seq revealed altered neurodevelopmental gene expression at the NPC stage and during MN differentiation, together with increased expression of stress- and apoptosis-associated genes. Proteomic analysis identified changes in proteins involved in neuronal structure, synaptic function, RNA regulation, and cellular homeostasis. Concordant decreases in selected RNAs and proteins supported disruption of neuronal structural and synaptic programs. Together, these findings reveal a stage-dependent requirement for SETX and identify SETX as an important regulator of transcriptional programs governing human MN maturation, function, and survival, providing a mechanistic framework for understanding the selective MN vulnerability associated with SETX-linked neurological diseases.

## INTRODUCTION

Senataxin (SETX) is a large nuclear protein (∼303 kDa) encoded by the *SETX* gene on human chromosome 9q34.13. It contains an N-terminal protein-interaction region and a conserved C-terminal superfamily-1 DNA/RNA helicase domain [1]. SETX is the human ortholog of yeast Sen1 and contributes to transcription, RNA processing, and genome stability [2–5]. It is expressed in many tissues, including the brain, spinal cord, and testis [6, 7]. SETX is found mainly in the nucleus and can accumulate at sites of DNA damage [8]. Biochemical studies have shown that SETX can unwind the RNA hybrids within R-loops and promote transcription termination [9, 10]. Other studies support roles in transcription elongation, replication restart, and the resolution of transcription-replication conflicts [4, 11, 12]. These functions link RNA regulation to genome maintenance and may help support the long-term function of neurons.

Mutations in *SETX* cause distinct inherited neurological disorders, highlighting its importance in the nervous system. Heterozygous missense mutations cause amyotrophic lateral sclerosis type 4 (ALS4), a rare, autosomal dominant motor neuron (MN) disease. ALS4 usually begins in childhood or adolescence and progresses slowly. Its main features include distal muscle weakness and atrophy and pyramidal signs [13, 14]. In contrast, biallelic loss-of-function mutations cause ataxia with oculomotor apraxia type 2 (AOA2). This autosomal recessive disorder is characterized by progressive cerebellar ataxia, peripheral axonal neuropathy, and elevated serum α-fetoprotein [1]. These distinct clinical features suggest that different forms of SETX dysfunction affect the nervous system in different ways.

SETX dysfunction may also contribute to MN degeneration in disorders without *SETX* mutations. In experimental models of spinal muscular atrophy (SMA), reduced survival motor neuron (SMN) protein was associated with lower SETX levels, R-loop accumulation, and increased DNA double-strand breaks. Restoring SETX reduced R-loops and DNA damage and improved neuronal survival [15]. Together with the MN involvement in ALS4, these findings suggest that SETX helps maintain MN function and survival.

MN development involves a series of coordinated steps, from proliferating neural progenitor cells (NPCs) to postmitotic neurons and functionally mature MNs. In the developing spinal cord, sonic hedgehog and retinoic acid signaling help establish MN progenitors that express OLIG2 [16]. Transcription factors such as NEUROG2, ISL1/2, LHX3, and MNX1/HB9 then help guide MN specification and differentiation [16–18]. Further maturation requires axonal and dendritic growth, expression of ion channels and synaptic proteins, and development of electrical activity and synaptic connections. These changes depend on coordinated gene expression and RNA regulation [19, 20]. Given its roles in these processes, SETX may support MN differentiation and maturation as well as neuronal survival.

Despite advances in SETX biology, its roles across stages of human MN development remain incompletely understood. Important mechanistic insights have come from studies of the yeast ortholog Sen1 [21, 22], immortalized human cell lines [10], patient fibroblasts [23], and animal models [7, 24]. These studies have linked SETX dysfunction to altered RNA regulation, cellular stress, and neuronal degeneration. However, how complete SETX loss affects different stages of human MN development remains poorly understood. A key question is whether SETX deficiency mainly affects neuronal maintenance or also alters gene expression during differentiation and maturation. Examining these effects may help explain how SETX dysfunction contributes to MN vulnerability.

Here, we examined the role of SETX in human MN development, maturation, and survival using complementary loss-of-function models. We generated isogenic SETX knockout (KO) human induced pluripotent stem cell (hiPSC) lines and differentiated them through NPCs into MNs. We also used shRNA-mediated knockdown during hiPSC-derived MN differentiation and directly converted WT and SETX KO fibroblasts into MNs. We combined survival and morphological analyses with multielectrode array (MEA) recordings, RNA hybrid measurements, RNA-seq, and quantitative proteomics. SETX loss slowed proliferation in hiPSCs and NPCs and impaired neurite development, electrical activity, and survival in MNs. It also increased RNA hybrid accumulation and altered gene expression during the NPC-to-MN transition. Integrated RNA and protein analyses identified changes in neuronal development, synaptic function, and cellular stress responses. Together, these findings support a role for SETX across stages of human MN development, maturation, and maintenance. They provide cellular and molecular insights into MN vulnerability associated with SETX dysfunction.

## RESULTS

### Generation and characterization of SETX knockout hiPSC lines and isogenic controls

Senataxin has diverse cellular functions, and disease-associated SETX mutations can impair motor neuron (MN) function [14, 24]. To investigate the role of SETX in human MNs, we generated three SETX knockout (KO) hiPSC lines and corresponding wild-type (WT) isogenic controls from the healthy donor line WTC11 (**Table S1**). DNA sequencing confirmed frameshift mutations in SETX in all three KO clones (**Figure S1**).

All newly generated clones showed typical pluripotent stem cell morphology, including a high nucleus-to-cytoplasm ratio (**Figure 1A**). They retained a normal male karyotype (46, XY; **Figure S2A**) and short tandem repeat profiling matched the parental WTC11 line (**Figure S2B**). We assessed pluripotency using previously described methods [25–28]. Immunocytochemistry and/or RT-PCR detected robust expression of pluripotency markers, including OCT3/4, NANOG, SOX2, SSEA4, and KLF4 (**Figures 1B, 1C**, **and S2C**).

**Figure 1.**
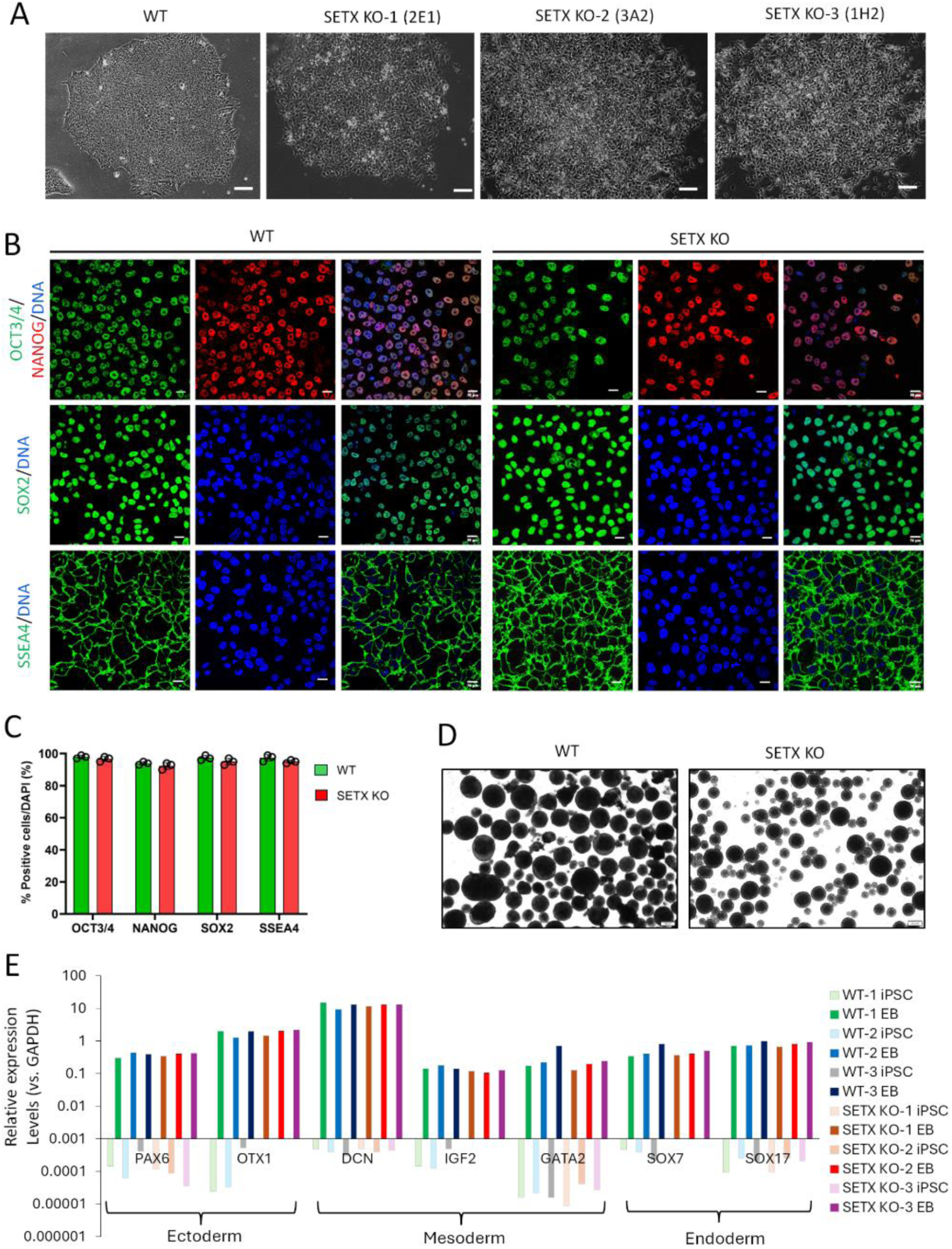
Generation and validation of SETX KO iPSC lines and isogenic controls. (**A**) Reprehensive micrographs showing the typical morphology of WT and SETX KO lines (three clones) with a high nucleus/cytoplasm ratio. Scale bars: 100 µm. (**B**) Representative confocal micrographs showing the expression and localization of indicated pluripotency markers. Scale bars: 50 µm. (**C**) Quantification of the percentage of cells (vs. DAPI) expressing pluripotent markers. N (clones) = 3 in each group. n (cell) = 100 for each clone. (**D**) Reprehensive micrographs showing embryoid bodies (EBs) derived from WT and SETX KO-2E1 iPSCs. Scale bars: 200 µm. (**E**) RT-PCR results showing the upregulation of three linage markers: ectoderm (PAX6, OTX1), mesoderm (DCN, IGF2, GATA2) and endoderm (SOX7, SOX17) in EBs compared to iPSCs.

All lines formed embryoid bodies (EBs) during spontaneous differentiation (**Figure 1D**). Compared with undifferentiated hiPSCs, EBs showed increased expression of markers of the three germ layers: ectoderm (PAX6 and OTX1), mesoderm (DCN, IGF2, and GATA2), and endoderm (SOX7 and SOX17). Expression levels were comparable to those in parental WTC11-derived EBs (**Figure 1E**). Together, these results show that the SETX KO hiPSC lines and WT isogenic control lines retained a normal karyotype and pluripotency-associated features, including the ability to express markers of all three germ layers after differentiation.

### SETX loss slows hiPSC and NPC proliferation and compromises MN survival

We generated hiPSC-derived MNs using our established differentiation protocol. hiPSCs were first differentiated into neural progenitor cells (NPCs), which were then induced to differentiate into MNs by lentiviral delivery of lineage-specific transcription factors (**Figure 2A**) [29, 30]. To examine the effects of SETX loss at different developmental stages, we assessed cell growth and survival in hiPSCs, NPCs, and differentiated MNs. Compared with WT controls, SETX KO hiPSCs formed less compact colonies and proliferated more slowly, with a significantly longer doubling time (**Figures 1A and 2B**). EBs derived from SETX KO hiPSCs were also significantly smaller than those derived from WT controls (**Figures 1D and 2C**). SETX KO hiPSCs generated NPCs, but these cells continued to proliferate significantly more slowly than WT NPCs (**Figures 2B and S3A**). These findings indicate that SETX loss reduces proliferation in hiPSCs and NPCs without preventing neural progenitor generation.

**Figure 2.**
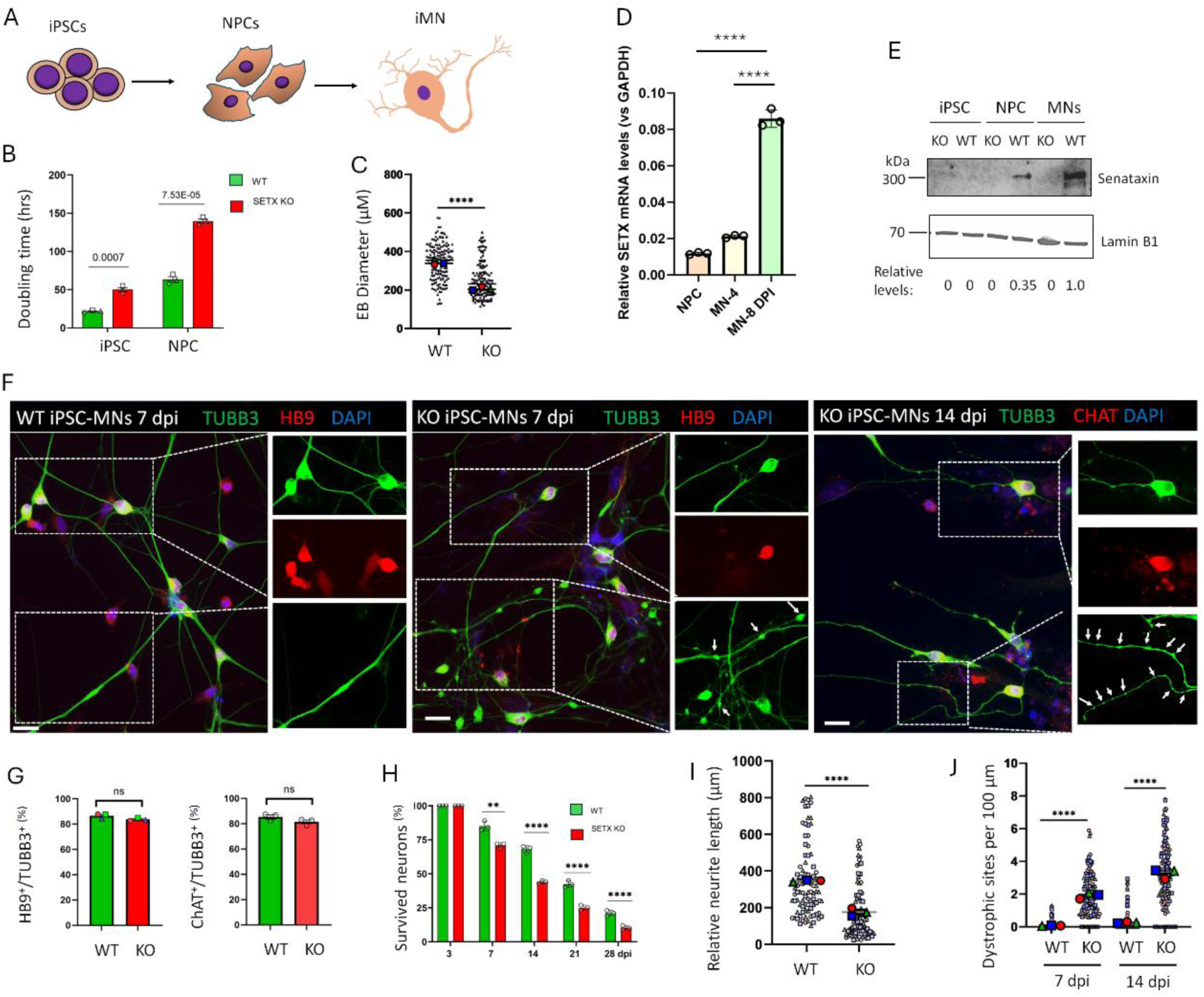
Differential effects of SETX KO on cell growth and survival during differentiation from hiPSC to motor neurons. (A) Schematic shows the three differnation stages from iPSC, NPC to MNs. (**B**) Doubling time of iPSC and NPC in WT and SETX KO lines. N = three independent experiments. (three clones), Student’s t-test. (**C**) The size of embryonic bodies (Ebs) derived from WT and SETX KO hiPSC lines. N = three clones. n (EB) = 80 for each clone. **** p<0.0001. Student’s t-test. (**D**) RT-PCR results showing the relative SETX mRNA levels in WT NPC, and induced MNs at 4 and 8 days. N = three clones (**E**) Western blotting of SETX protein in WT and SETX KO lines at indicated differentiation stages. Nuclear Lamin B1 serves as a loading control. The relative SETX protein levels normalized to Lamin B1 was indicated at the bottom. (**F**) Reprehensive micrographs showing MNs derived from WT and SETX KO hiPSC lines at 7 and 14 dpi with immunostaining of indicated markers. White dotted boxes were highlighted soma or neurites. Scale bars: 100 µm. (**G**) Quantification of the percentage of neurons with MN marker HB9 and CHAT. N (clones) = 3 in each group. n (cell) = 100 for each clone. Student’s t-test. (**H**) Quantification of survived MNs derived from WT and SETX KO lines at indicated time points. N (clones) = 3 in each group. n (neurons) > 500 for each clone at each time point. (**I**) Relative neurite length of MNs at 7 dpi. (**J**) Quantification of dystrophic sites normalized with neurite length under indicated conditions. N (clones) = 3 in each group. n (neurons) =100 for each clone in each condition.

Both WT and SETX KO cells differentiated into MNs. In WT cells, SETX mRNA and protein levels increased during differentiation from hiPSCs to NPCs and MNs (**Figures 2D and 2E**). When co-cultured with astrocytes [31], SETX KO MNs developed neuronal morphology and expressed the MN markers HB9 (MNX1) and choline acetyltransferase (ChAT), similar to WT controls (**Figures 2F and 2G**). However, SETX KO MNs showed progressive neuronal loss (**Figure 2H**), significantly shorter neurites (**Figure 2I**), and more dystrophic neurite swellings (**Figure 2J**). Thus, although SETX KO cells acquired MN markers, their subsequent survival and neurite integrity were impaired.

To further assess MN survival, we cultured MNs without astrocytes. Under these conditions, SETX KO MNs rapidly degenerated, with nearly all neurons dying by 5 days post-viral infection (DPI). In contrast, only a small fraction of WT MNs showed degeneration at the same time point (**Figures S3B–S3D**). Together, these results show that SETX loss slows proliferation in hiPSCs and NPCs and compromises the survival and morphological integrity of differentiated MNs. These findings support a role for SETX in MN maintenance.

### SETX KO reduces spontaneous firing and network activity in hiPSC-derived MNs

To assess the functional maturation of SETX KO MNs, we cultured WT and SETX KO MNs on multielectrode array (MEA) plates, sequentially seeding MNs and astrocytes onto the microelectrode grids with similar cell density (**Figure 3A**). With astrocyte support, a subset of SETX KO MNs survived for several weeks and developed mature neuronal morphology by 14 DPI, allowing longitudinal recordings [31]. WT MNs began to fire spontaneously at approximately 10 DPI. Spike amplitudes increased during maturation, and firing rates peaked around 20 DPI. In contrast, SETX KO MNs showed markedly reduced spontaneous firing throughout the recording period and significantly lower spike amplitudes than WT controls (**Figures 3B**, **3C, and Videos 1-4**). Raster plots also showed reduced burst activity in SETX KO cultures (**Figure 3D**). Single-electrode burst frequency was significantly lower than in WT cultures, whereas burst duration did not differ significantly (**Figures 3E and 3F**). Synchronized network bursts were almost absent in SETX KO cultures, indicating reduced coordination of neuronal activity across the electrode array (**Figures 3G–3I**). Together, these findings show that SETX loss reduces spontaneous firing, burst frequency, and synchronized network activity in human MN cultures, consistent with impaired functional maturation.

**Figure 3.**
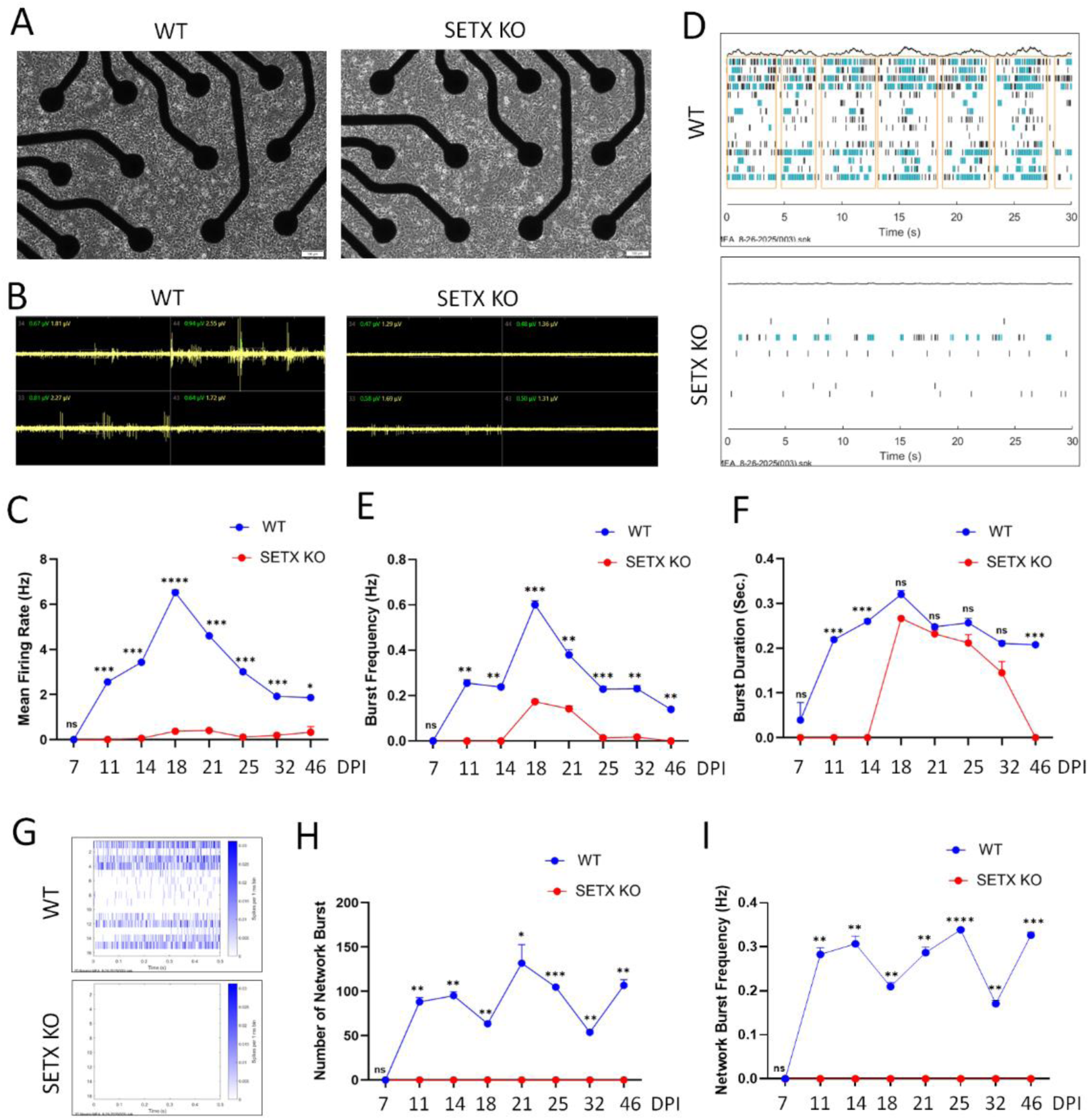
SETX KO impairs the electrophysiological maturation of hiPSC-derived MNs. **(A)** Representative micrographs of hiPSC-derived MNs cultured on multielectrode array (MEA) plates at 7 days post-viral infection (DPI). Scale bars, 100 µm. **(B)** Representative continuous extracellular waveform recordings from individual electrodes at 21 dpi. **(C)** Mean firing rate of WT and SETX KO MNs at the indicated time points from 7 to 46 dpi. **(D)** Representative raster plots of spontaneous neuronal activity recorded across the 16-electrode array at 21 dpi. Individual detected spike events are shown in black, and cyan-shaded regions indicate detected burst activity. **(E)** Quantification of burst frequency at the indicated time points. **(F)** Quantification of burst duration at the indicated time points. **(G)** Representative network-burst activity recorded across the 16-electrode array at 21 dpi. **(H)** Quantification of the number of network bursts at the indicated time points from 7 to 46 dpi. **(I)** Quantification of network-burst frequency at the indicated time points from 7 to 46 dpi. For **(C–F, H, I)**, data are presented as mean ± SEM from three independent experiment. In each experiment, three MEA wells per genotype and three 5-min recordings were acquired from each well at each time point. Statistical significance between WT and SETX KO groups at each time point was determined using multiple two-sided unpaired *t*-tests with Welch’s correction. ns, not significant; *P < 0.01; **P < 0.001; ***P < 0.0001; ****P < 0.00001.

### SETX downregulation compromises MN survival and neurite integrity

To further assess the role of SETX in human MN development, we generated lentiviral vectors expressing SETX-targeting shRNAs together with an mCherry reporter. Each shRNA targeted a distinct region of the SETX coding sequence and reduced SETX expression compared with a control vector expressing mCherry alone (**Figure 4A**). WT NPCs transduced with SETX shRNAs differentiated into MNs with typical neuronal morphology (**Figure 4B**). During the first 5 DPI, SETX knockdown (KD) MNs showed neurite outgrowth, morphology, and survival comparable to those of control MNs (**Figure 4C**). However, after 7 DPI, SETX KD MNs rapidly degenerated, with most neurons dying by approximately 11 DPI (**Figure 4D**). Surviving SETX KD MNs also had significantly shorter neurites than control MNs (**Figure 4E**). These findings support a role for SETX in maintaining MN survival and neurite integrity. In cultures without astrocytes, most SETX KD MNs died by approximately 11 DPI, whereas nearly all SETX KO MNs died by 5 DPI (**Figures S3 and 4D**). This later onset of extensive neuronal loss following SETX KD is consistent with the possibility that residual SETX expression delays degeneration.

**Figure 4.**
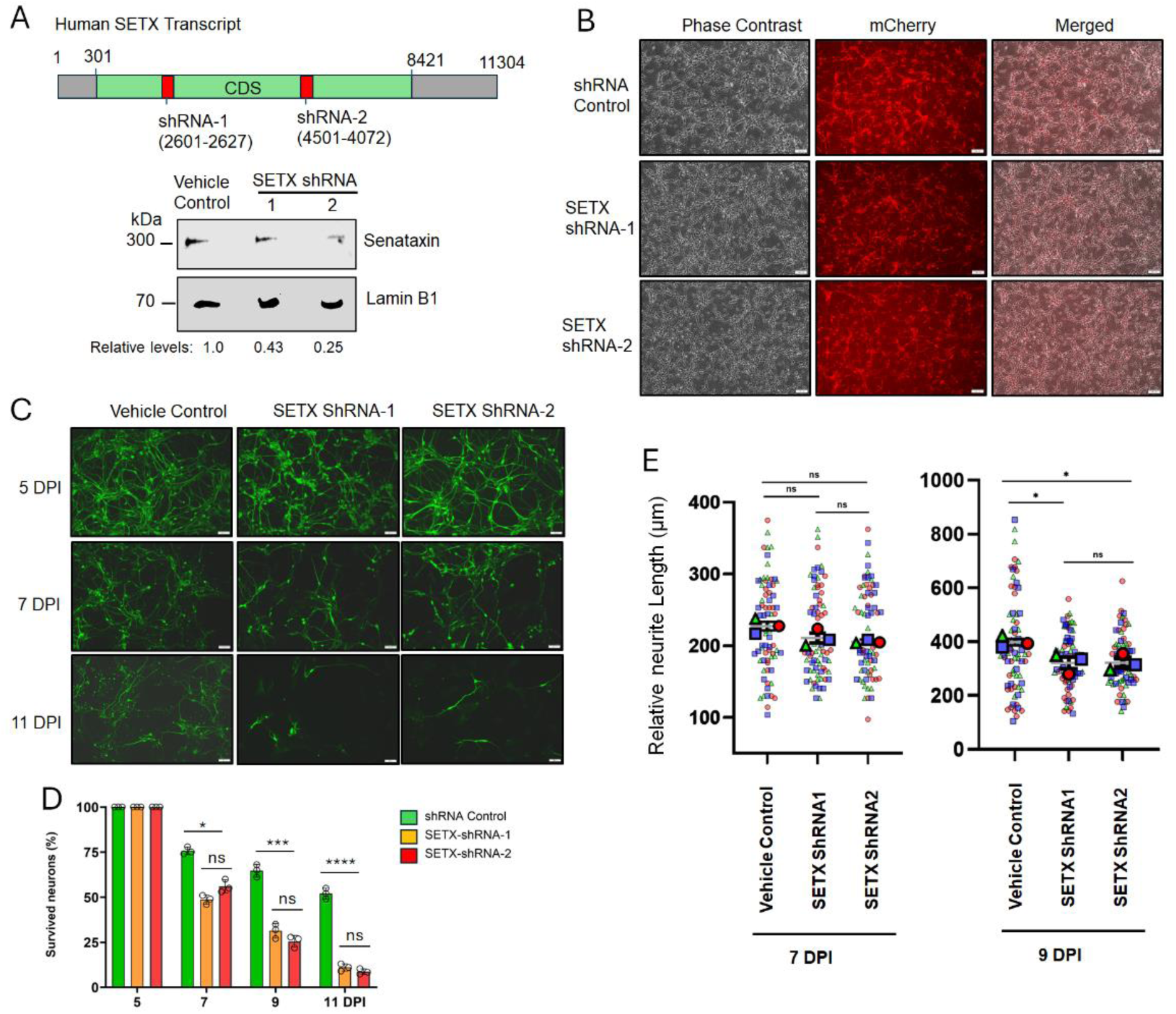
SETX downregulation compromises motor neuron maturation and survival. **(A)** Schematic illustrating two shRNAs targeting distinct regions of the *SETX* coding sequence (top) and Western blot analysis confirming efficient downregulation of SETX protein in hiPSC-derived MNs (bottom). **(B)** Representative fluorescence micrographs showing WT hiPSC-derived MNs expressing SETX-targeting shRNAs (mCherry-positive) at 5 DPI. Scale bars, 100 µm. **(C)** Representative micrographs showing hiPSC-derived MNs under the indicated conditions at 5, 7, and 11 DPI, visualized by TUBB3 immunostaining. Scale bars, 100 µm. **(D)** Quantification of surviving MNs under the indicated conditions at different time points. N = 3 independent experiments; n > 100 neurons per experiment at each time point. **(E)** Quantification of relative neurite length of hiPSC-derived MNs under the indicated conditions. N = 3 independent experiments; n > 100 neurons per experiment per group. Individual experiments are indicated by distinct colors. For **(D, E)**, statistical significance was determined by one-way ANOVA followed by Dunnett’s multiple-comparisons test. ns, not significant; *P < 0.05; ***P < 0.001; ****P < 0.0001.

### Directly converted MNs from SETX KO fibroblasts show impaired morphological maturation

To examine whether SETX loss also affects MN maturation in a second model, we directly converted human fibroblasts into MNs (diMNs; **Figure 5A**). We generated a SETX KO line from immortalized human fibroblasts (**Figures 5B, S1D, and Table S1**). Compared with the WT isogenic control, SETX KO fibroblasts proliferated more slowly but showed no obvious differences in morphology or viability. Using our established protocol [32], we converted WT and SETX KO fibroblasts into diMNs. Conversion efficiencies were comparable between genotypes, as assessed by expression of the MN markers HB9 and ChAT (**Figures 5C–5E**). Despite acquiring MN markers, SETX KO diMNs developed abnormal neuronal morphology. WT diMNs typically extended a single long axon-like process, whereas most SETX KO diMNs formed multiple short, thin neurites (**Figures 5C and 5F**). SETX KO diMNs also had significantly smaller soma area (**Figure 5G**) and markedly shorter neurites (**Figure 5H**). Between 14 and 21 DPI, a period of diMN maturation [33, 34], neurite length in SETX KO diMNs showed little further increase, whereas WT neurites continued to elongate (**Figure 5H**). SETX KO diMNs also developed more dystrophic neurite swellings than WT controls (**Figures 5C and 5I**). Together, these findings indicate that SETX loss impairs morphological maturation and neurite integrity in directly converted human MNs.

**Figure 5.**
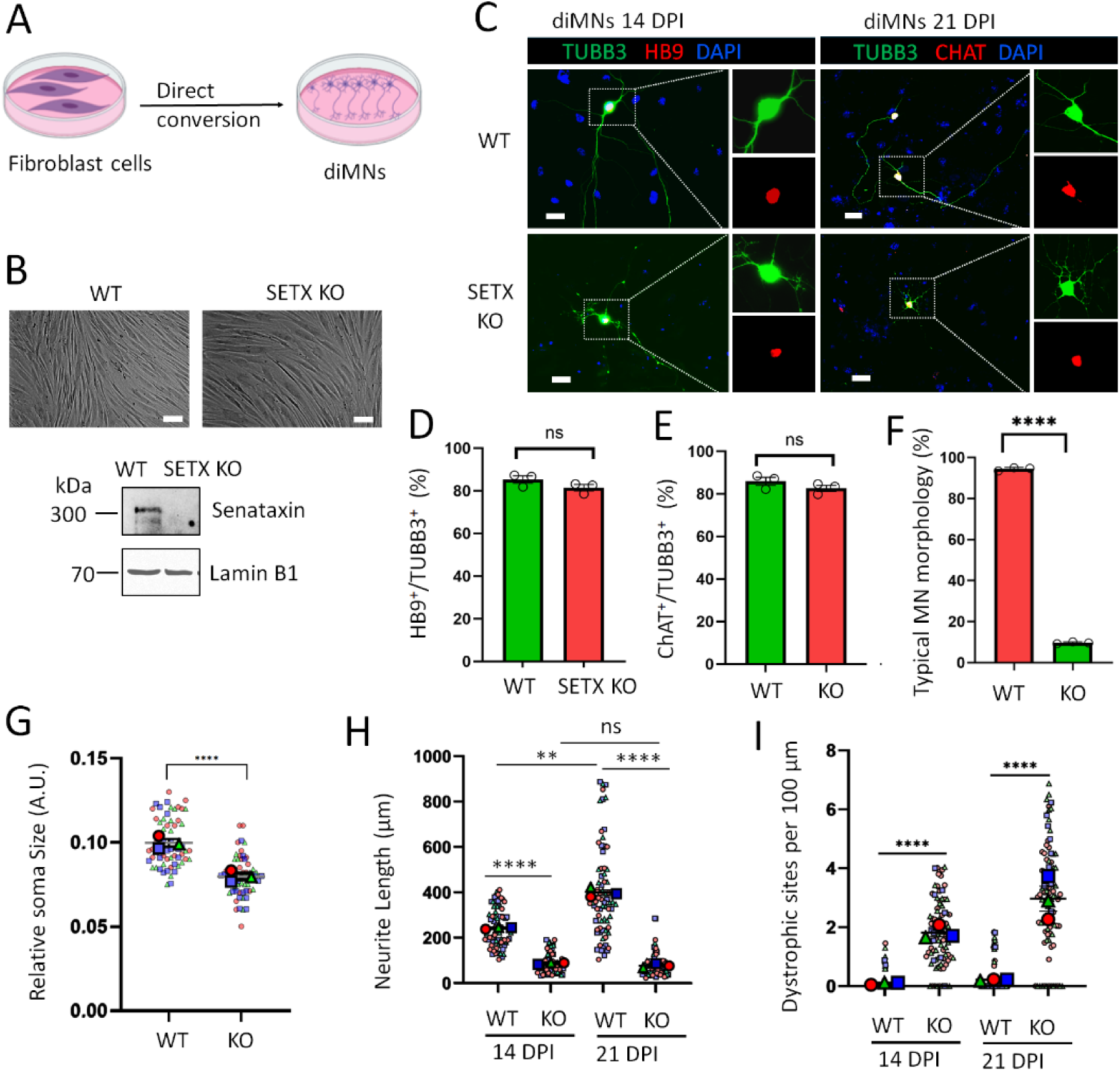
Directly converted MNs from SETX KO fibroblasts exhibit impaired maturation. (**A**) Schematic showing direct conversion fibroblasts to motor neurons. (**B**) Micrographs of WT and SETX KO fibroblasts and Senataxin levels measured by western blotting. Nuclear Lamin B1 serves as a loading control. Scale bars = 100 µm. (**C**) Immunocytochemistry of motor neuron markers HB9 and CHAT in diMNs at 14 and 21 dpi, respectively. Scale bars = 50 µm. (**D**) Percentage of HB9 (+) cells per TUBB3 (+) cells at 14 DPI. (**E**) Percentaz ge of CHAT (+) cells per TUBB3 (+) cells at 21 DPI. (**F**) Percentage neurons with typical MN morphology at 14 dpi. (**G**) Relative soma size (area) of diMNs at 14 dpi. (**H**) Relative neurite length of diMNs at 14 and 21 dpi. **(I)** Quantification of dystrophic sites normalized to neurite length of diMNs at 14 and 21 dpi. For **(D–I)**, n > 50 neurons from three independent experiments. ns, not significant; ** P<0.01, ****P < 0.0001, Student’s *t*-test.

### SETX loss increases RNA hybrid accumulation with the largest relative increase in human MNs

SETX contributes to R-loop resolution during transcription in multiple model systems [9–11]. To compare the effects of SETX loss across cell types, we assessed RNA hybrids using the S9.6 monoclonal antibody [35, 36]. Immunocytochemistry showed significantly higher S9.6 signal intensity in SETX KO fibroblasts (**Figures S4A**), NPCs (**Figures S4C**), and hiPSC-derived MNs (**Figure S4D**) than in their respective WT controls. RNase H treatment reduced the signal, supporting the detection of RNA hybrids (**Figure S4B**). The KO-to-WT ratio of nuclear S9.6 signal intensity was significantly higher in MNs than in fibroblasts or NPCs (**Figure S4E**).

We further assessed hybrid accumulation using S9.6 dot-blot analysis [37, 38]. RNase H treatment reduced the dot-blot signal (**Figure S4F**). After normalization to dsDNA loading, SETX KO NPCs and MNs showed higher hybrid signals than their respective WT controls (**Figures S4G–S4J**). The KO-to-WT ratio was significantly higher in MNs than in NPCs (**Figure S4K**). Together, these results support increased R-loop accumulation following SETX loss, with the largest relative increase in human MNs among the cell types examined.

### Bulk RNA sequencing reveals altered neuronal development and stress-response programs in SETX KO MNs

To investigate the molecular changes associated with impaired MN development and maturation following SETX loss, we performed bulk RNA sequencing (RNA-seq) on purified hiPSC-derived MNs at 4 DPI, at which the viability and neuronal purity were comparable between SETX KO MNs and isogenic WT controls (**Figure 6A**). We identified 3,976 differentially expressed genes (DEGs) among 24,510 detected transcripts. These included 1,854 downregulated and 2,122 upregulated genes (**Figure 6B and Table S4**).

**Figure 6.**
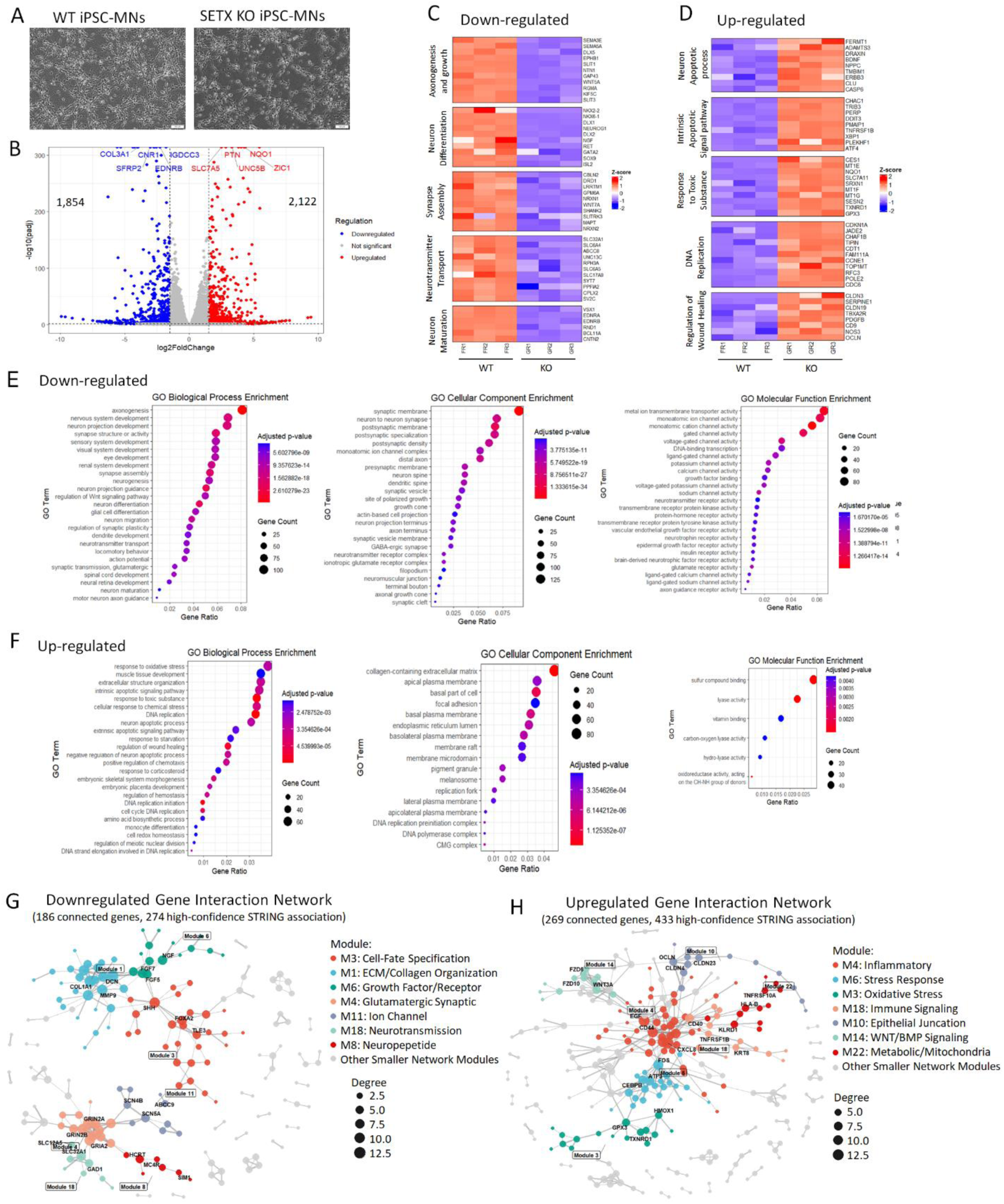
Bulk RNA sequencing reveals widespread disruption of neuronal function and cellular homeostasis in SETX KO MNs. (A) Representative micrographs of WT and SETX KO hiPSC-derived MN cultures at 4 DPI, prior to collection for RNA isolation. Scale bars, 100 µm. (**B**) Volcano plot showing differentially expressed genes (DEGs) in SETX KO MNs relative to WT MNs. Genes with an adjusted *P* value < 0.05 and an absolute fold change ≥ 1.5 (|log2FC| ≥ 0.585) were considered differentially expressed. (**C, D**) Heatmaps showing selected significantly downregulated (C) and upregulated (D) genes associated with biological processes relevant to neuronal function and cellular homeostasis. Heatmaps show row-wise Z scores of normalized gene expression across three independent biological replicates per genotype, with red and blue indicating relatively higher and lower expression, respectively. (**E, F**) Gene Ontology (GO) enrichment analysis of significantly downregulated (E) and upregulated (F) genes in SETX KO MNs, respectively. Enriched terms are shown for Biological Process (BP), Cellular Component (CC), and Molecular Function (MF) categories. (**G, H**) Network visualization of downregulated genes (G) and upregulated genes (H) with different modules highlighted by distinct colors.

Gene Ontology (GO) analysis showed that downregulated genes were enriched in processes related to neuronal development and maturation. These included axonogenesis, neurite outgrowth, neuronal differentiation, synapse assembly, neurotransmitter transport, and neuron maturation (**Figures 6C and 6E and Table S4).** Downregulated genes included the axon guidance genes SEMA3E, SEMA5A, EPHB1, SLIT1, and NTN1, as well as developmental transcription factors such as NKX2-2, NKX6-1, DLX1, DLX2, NEUROG1, NEUROG3, GATA2, SOX9, and ISL2.

Genes associated with synapse formation and plasticity, including CBLN2, LRRTM1, GPM6A, NRXN1, WNT7A, and SHANK2, were also downregulated. Additional decreases affected genes involved in synaptic function and neuronal signaling, including SLC32A1, UNC13C, RPH3A, SYT7, SYN3, CPLX2, SV2C, RET, and CNTN2. Network analysis grouped downregulated genes into modules associated with extracellular matrix and collagen organization (M1; COL1A1, COL3A1, BGN, and DCN) and developmental patterning (M7; SHH and FOXA2). Other modules were associated with synaptic signaling, ion channel activity, receptor signaling, neurotransmission, and neuroendocrine functions (M10, M4, M17, M5, and M3; **Figure 6G).** These expression changes are consistent with the neurite abnormalities, reduced electrophysiological activity, and neuronal loss observed in SETX KO MNs.

Upregulated genes were enriched in GO terms related to neuronal apoptosis, intrinsic apoptotic signaling, responses to toxic stimuli, DNA replication, and wound responses (**Figures 6D and 6F and Table S4**). Stress-and apoptosis-associated genes, including CHAC1, TRIB3, PERP, DDIT3, PMAIP1, CDKN1A, and TNFRSF1B, showed increased expression. This pattern is consistent with increased cellular stress, including endoplasmic reticulum (ER) stress, and altered apoptotic signaling. BDNF, NPPC, and ERBB3 were also upregulated, potentially reflecting compensatory changes in growth and survival signaling. Other upregulated genes were associated with detoxification (MT1E, MT1F, MT1X, and GSTO2), antioxidant defense (NQO1, SLC7A11, SRXN1, SESN2, TXNRD1, GPX3, and GSR), and extracellular matrix organization and cell adhesion (OPTC, SPOCK2, ITGA8, ITGA2, COL13A1, MMP25, and LAMA2). Genes associated with cell-cycle regulation, chromatin regulation, and DNA replication were also upregulated, including CDKN1A, JADE2, CHAF1B, TIPIN, CDT1, and RFC3. Network analysis identified modules related to oxidative and redox metabolism (M7), stress responses (M9), inflammatory and immune signaling (M1 and M17), cell adhesion and signaling (M8), and WNT signaling (M3; **Figure 6H**). Together, these findings show that SETX loss alters gene expression programs associated with neuronal development and synaptic function while increasing the expression of stress-response genes. These changes may contribute to impaired MN maturation and survival.

### SETX loss alters gene expression programs during the NPC-to-MN transition

To examine the effects of SETX loss at the progenitor stage, we performed RNA-seq on WT and SETX KO NPCs. We then compared changes in gene expression during MN differentiation between the two genotypes. SETX KO NPCs exhibited 4,043 DEGs relative to WT NPCs, including 1,635 downregulated and 2,408 upregulated genes (**Figure S5A and Table S5**). Downregulated genes were enriched in GO terms related to neurogenesis, nervous system development, neuronal fate commitment, transcriptional regulation, and ion channel activity (**Figures S5B–S5D and Table S5**). Other enriched processes included spinal cord development, visual system development, and eye morphogenesis. Upregulated genes were enriched in cell-substrate adhesion, extracellular structure organization, wound responses, regionalization, and hemostasis (**Figures S5E and S5F**). These findings indicate that SETX loss alters neurodevelopmental gene expression programs at the NPC stage.

We next examined changes in gene expression during the NPC-to-MN transition. In WT cells, 2,277 genes were downregulated and 2,809 were upregulated in MNs relative to NPCs (**Figure S6A and Table S6**). Downregulated genes were enriched in cell-cycle regulation, DNA replication, and cell division (**Figures S6C and S6E and Table S6**). Upregulated genes were enriched in neuronal development and maturation processes, including axonogenesis, neuron projection guidance, synapse assembly, and neurotransmitter transport (**Figures S6D and S6F and Table S6**). Thus, WT differentiation involved coordinated suppression of proliferative programs and increased expression of genes associated with neuronal maturation.

SETX KO cells also showed broad changes during differentiation, with 2,344 downregulated and 2,491 upregulated genes (**Figure S6B and Table S7**). However, the sets of genes meeting the differential expression criteria differed between genotypes. Among genes upregulated during differentiation, 1,122 met these criteria only in WT cells (Up WT Only), 804 only in SETX KO cells (Up KO Only), and 1,687 in both genotypes (**Figure 7A**). Among shared upregulated genes, subsets showed greater (More Up) or weaker (Less Up) induction in SETX KO cells than in WT cells (**Figure 7B and Table S7**).

**Figure 7.**
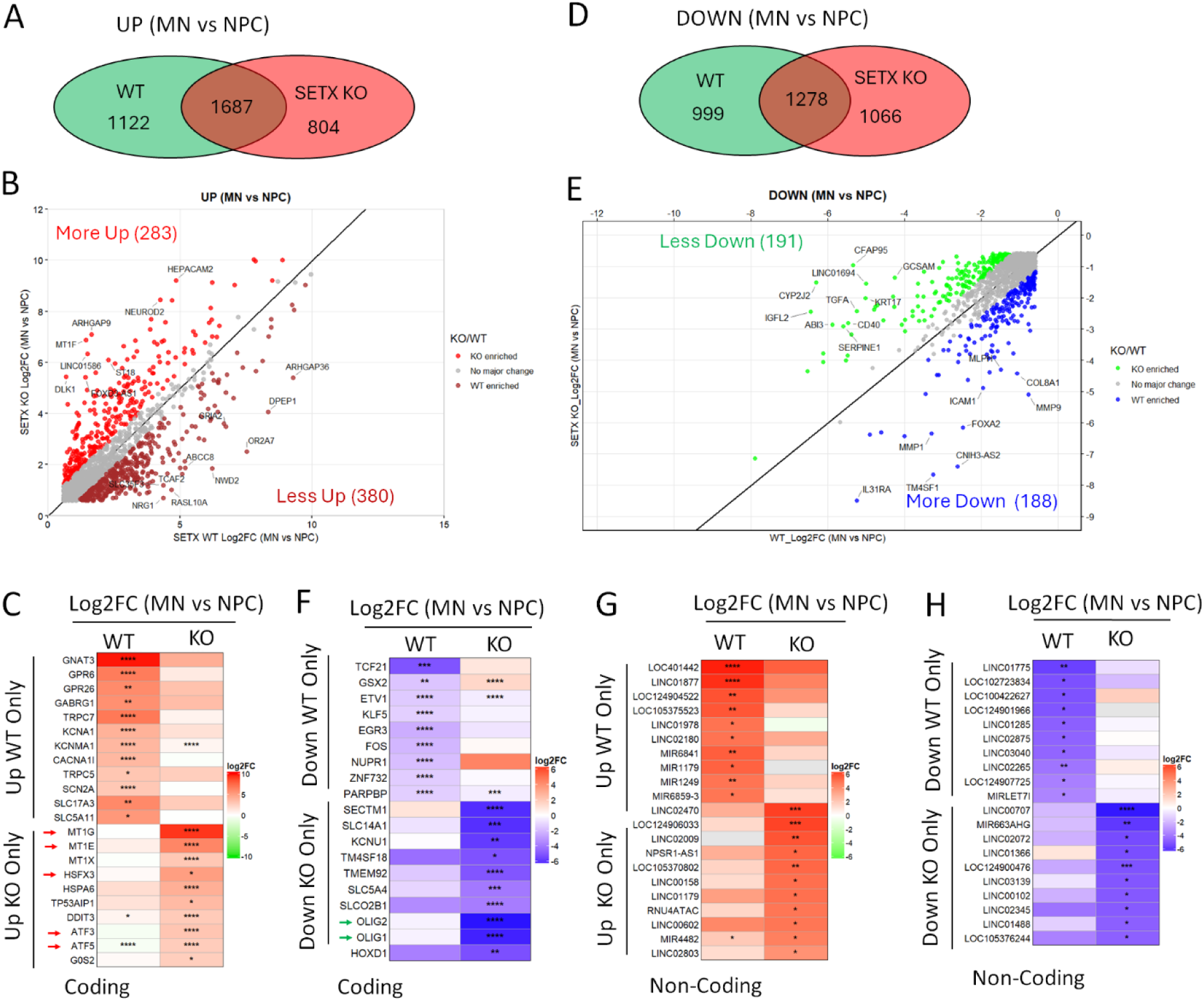
SETX loss disrupts the transcriptional transition from neural progenitor cells to motor neurons. (**A**) Venn diagram showing the numbers of genes significantly upregulated during the transition from neural progenitor cells (NPCs) to motor neurons (MNs) in WT and SETX KO cells, including genes uniquely upregulated in each genotype and those shared between genotypes. (**B**) Comparison of expression changes in shared genes that were significantly upregulated during the NPC-to-MN transition in both WT and SETX KO cells. (**C**) Heatmap showing selected protein-coding genes with differential upregulation during the NPC-to-MN transition in WT and SETX KO cells. (**D**) Venn diagram showing the numbers of genes significantly downregulated during the transition from NPCs to MNs in WT and SETX KO cells, including genes uniquely downregulated in each genotype and those shared between genotypes. (**E**) Comparison of expression changes in shared genes that were significantly downregulated during the NPC-to-MN transition in both WT and SETX KO cells. (**F**) Heatmap showing selected protein-coding genes with differential downregulation during the NPC-to-MN transition in WT and SETX KO cells. (**G**) Heatmap showing selected non-coding RNAs with differential upregulation during the NPC-to-MN transition in WT and SETX KO cells. (**H**) Heatmap showing selected non-coding RNAs with differential downregulation during the NPC-to-MN transition in WT and SETX KO cells. For (C, F–H), *P* < 0.05; P < 0.01; *P* < 0.001; P < 0.0001.

Genes upregulated only in WT cells included genes associated with neuronal signaling and function. These encoded G proteins and receptors (GNAT3, GPR6, GPR26, and GABRG1), ion channels (TRPC7, KCNA1, CACNA1I, and SCN2A), and solute carriers (SLC17A3 and SLC5A11; **Figure 7C**). Genes upregulated only in SETX KO cells included several stress-associated genes. These included metallothioneins (MT1G, MT1E, and MT1X), heat-shock-related genes (HSFX3 and HSPA6), and genes associated with stress and apoptosis (TP53AIP1, DDIT3, G0S2, ATF3, and ATF5) [39–43]. These divergent transcriptional programs indicate that SETX KO NPCs fail to appropriately activate a subset of neuronal maturation programs while exhibiting enhanced activation of cellular stress pathways during the transition to MNs. These molecular changes are consistent with the impaired neuronal maturation and premature degeneration observed in both hiPSC-derived and directly converted MNs.

The sets of downregulated genes also differed between genotypes. During differentiation, 999 genes met the downregulation criteria only in WT cells (Down WT Only), 1,066 only in SETX KO cells (Down KO Only), and 1,278 in both genotypes (**Figure 7D and Table S7**). Shared downregulated genes included subsets with greater (More Down) or weaker (Less Down) downregulation in SETX KO cells (**Figure 7E**). Genes downregulated only in WT cells included transcriptional regulators such as TCF21, FOXQ1, GSX2, ETV1, and KLF5, as well as NUPR1, ZNF732, and PARPBP (**Figure 7F**). Genes downregulated only in SETX KO cells included OLIG1 and OLIG2 and genes encoding membrane-associated proteins, including SECTM1, SLC14A1, KCNU1, TM4SF18, and TMEM92 (**Figure 7F**). OLIG1 and OLIG2 are involved in neural development, and OLIG2 has an established role in spinal MN specification [16, 44, 45]. Their expression changes identify OLIG-associated developmental programs as candidates for further investigation.

To examine the biological processes represented by these DEG subgroups, we performed GO enrichment and gene interaction network analyses (**Figures S7 and S8**). Genes upregulated only in WT cells or showing weaker induction in SETX KO cells were enriched in neuronal maturation and function, including synaptogenesis and neurotransmitter regulation (**Figure S7A**). Corresponding network modules were associated with G protein-coupled receptors, ion channels, synaptic vesicle trafficking, and cell-surface receptors (**Figure S8A**). Genes upregulated only in SETX KO cells or showing greater induction in SETX KO cells were enriched in neuronal specification and differentiation, cell migration, and transcriptional regulation (**Figure S7B**). Network analysis identified modules associated with neuronal differentiation, amino acid transport, chromatin and transcriptional regulation, and cellular stress responses (**Figure S8B**). Thus, the KO-associated changes involved both developmental and stress-related programs.

Genes downregulated only in WT cells or showing weaker downregulation in SETX KO cells were enriched in mitosis, DNA replication, and cell-cycle regulation (**Figure S7C**). Their network modules were associated with cell-fate and stress signaling, redox metabolism, and DNA replication and repair (**Figure S8C**). Genes downregulated only in SETX KO cells or showing greater downregulation in SETX KO cells were enriched in extracellular matrix organization, cell migration, and cell adhesion (**Figure S7D**). Corresponding network modules were associated with inflammatory and antiviral responses, lipid metabolism, and transcriptional regulation (**Figure S8D**). These analyses indicate that SETX loss affects multiple gene expression programs during MN differentiation.

We also identified non-coding RNA transcripts whose expression patterns differed during WT and SETX KO differentiation (**Figures 7G and 7H**). Although the roles of many of these transcripts in MN development remain unclear, these findings suggest that SETX loss affects both coding and non-coding components of the differentiation-associated expression program. Together, these analyses show that SETX loss alters gene expression at the NPC stage and during the transition to MNs. The changes involve neuronal maturation, developmental regulation, proliferative programs, and cellular stress responses. These findings provide a molecular context for the impaired maturation, reduced electrophysiological activity, and neuronal loss observed in SETX-deficient human MNs.

### Proteomic profiling reveals altered neuronal maturation and homeostasis-associated networks in SETX KO MNs

To examine the effects of SETX loss at the protein level, we performed quantitative proteomic analysis of SETX KO and WT hiPSC-derived MNs at 4 DPI. We detected 6,755 proteins, of which 693 were differentially expressed using an adjusted *P* value < 0.05 and |log₂FC| ≥ 0.585 as thresholds. These included 438 downregulated and 255 upregulated proteins (**Figure 8A and Table S9**). GAPDH and the neuronal marker RBFOX3 (NeuN) did not differ significantly between genotypes. Differentially expressed proteins (DEPs) were associated with neuronal development, cell adhesion, synaptic function, RNA regulation, proteostasis, and mitochondrial homeostasis (**Figure 8B**).

**Figure 8.**
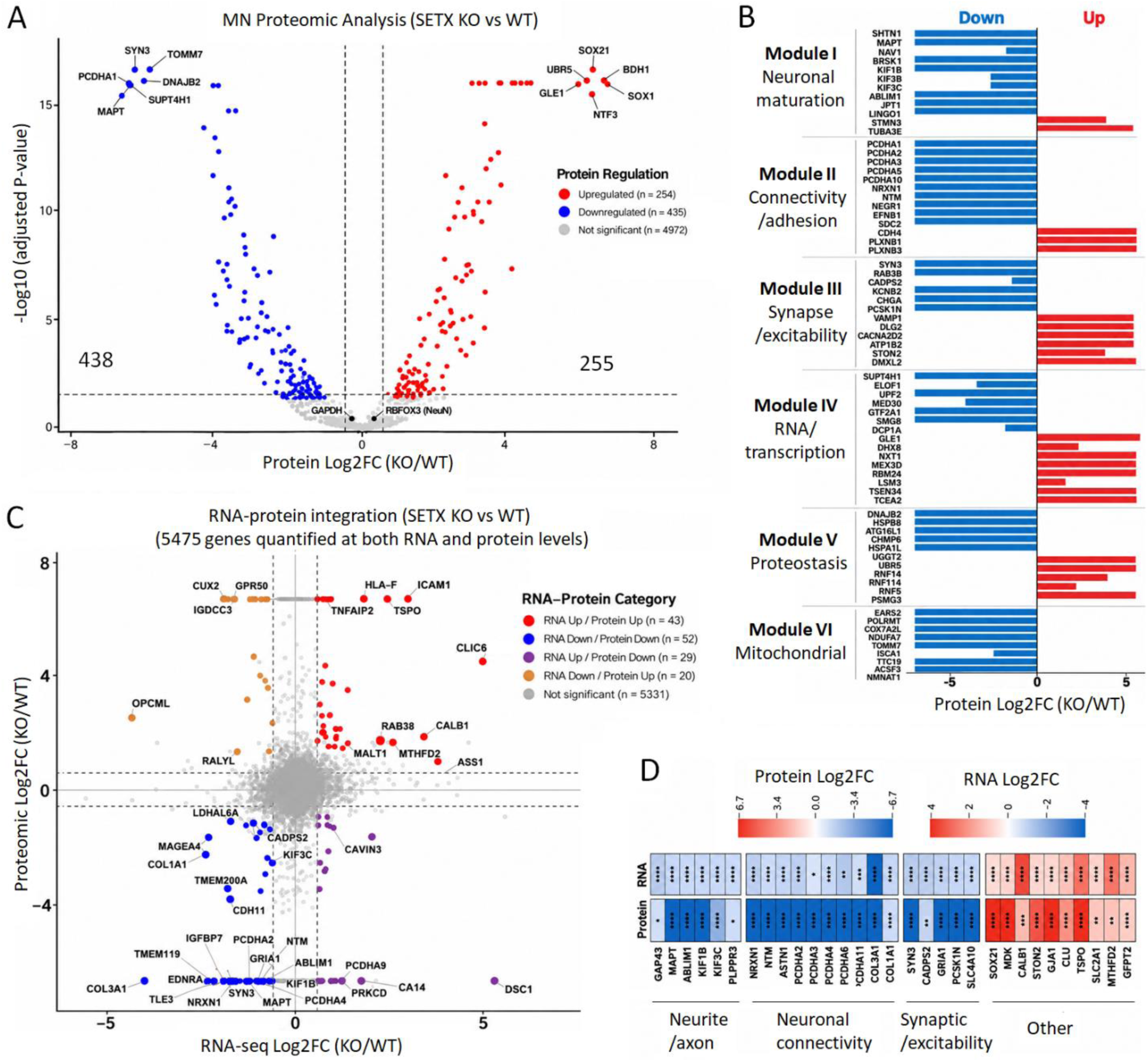
Proteomic profiling reveals impaired neuronal maturation and homeostatic networks in SETX KO motor neurons. (**A**) Volcano plot showing differentially abundant proteins in SETX KO MNs compared with WT MNs at 4 DPI. Proteins with an adjusted *P* value < 0.05 and an absolute fold change ≥ 1.5 (|log2FC| ≥ 0.585) were considered differentially expressed. (**B**) Selected proteins significantly downregulated or upregulated in SETX KO MNs, grouped into functional modules based on their molecular functions and biological roles. (**C**) Integrated transcriptomic and proteomic analysis showing four categories of RNA-protein expression changes based on the direction and concordance of changes at the mRNA and protein levels, including RNA Up/Protein Up (n=43), RNA Down/Protein Down (n=52), RNA Up/Protein Down (n=29), and RNA Down/Protein Up (n=20). (**D**) Heatmap showing selected genes exhibiting concordant changes at both the mRNA and protein levels in SETX KO MNs compared with WT MNs. * *P* < 0.05; ** P < 0.01; *** *P* < 0.001; **** P < 0.0001.

SETX KO MNs showed reduced abundance of proteins involved in neuronal differentiation, cytoskeletal organization, neurite development, and intracellular transport. These included SHTN1, MAPT, NAV1, ABLIM1, KATNB1, BRSK1, KIF1B, KIF3B, and KIF3C (Module I, **Figure 8B**). Several neuronal cell-surface proteins, including NEGR1, NTM, ASTN1, and LINGO1, were also reduced. Multiple members of the clustered protocadherin-alpha family decreased, along with other adhesion and connectivity proteins, including NRXN1, EFNB1, EFNB3, and SDC2. Selected cytoskeletal and cell-surface proteins, including STMN3, TUBA3E, CDH4, PLXNB1, and PLXNB3, increased (Module II, **Figure 8B**). Together, these changes indicate altered abundance of proteins associated with neuronal morphology, transport, and connectivity.

Proteins associated with synaptic transmission and membrane excitability also showed changes in both directions. SYN3, RAB3B, CADPS2, CHGA, PCSK1N, and the voltage-gated potassium channel KCNB2 were reduced. In contrast, VAMP1, DLG2, CACNA2D2, ATP1B2, STON2, and DMXL2 were increased (Module III, **Figure 8B**). These findings indicate broad changes in synaptic, secretory, and excitability-associated proteins. Such changes may contribute to the reduced firing, burst activity, and network synchronization observed in SETX KO MN cultures.

SETX loss also affected proteins involved in transcription and RNA regulation. Transcription-associated proteins, including SUPT4H1, ELOF1, POLR2L, MED8, MED30, GTF2A1/GTF2A2, TAF6, and SP1, were decreased. Components of RNA surveillance and turnover, including UPF2, SMG8, and DCP1A, were also reduced. Other proteins associated with RNA processing, transport, and regulation were increased, including GLE1, NXT1, DHX8, MEX3D, RBM24, RBM38, LSM3, TSEN34, and TCEA2 (Module IV, **Figure 8B**). These findings indicate that SETX loss affects multiple components of gene expression regulation.

Proteins associated with cellular homeostasis were also altered. Several components of protein quality control and degradation pathways, including DNAJB2, HSPB8, HSPA1L, ATG16L1, UGGT2, and CHMP6, were reduced. Ubiquitin- and proteasome-associated proteins, including UBR5, RNF14, RNF114, RNF5, and PSMG3, were increased (Module V, **Figure 8B**). Reduced mitochondrial proteins included EARS2 and POLRMT, which participate in mitochondrial gene expression, and the respiratory-chain-associated proteins NDUFA7, NDUFB2, COX7A2, and COX7A2L. TOMM7, ISCA1, TTC19, and ACSF3 were also reduced (Module VI, **Figure 8B**), indicating changes in proteins associated with mitochondrial import, iron-sulfur cluster biology, respiratory function, and metabolism. The NAD biosynthesis enzyme NMNAT1 was decreased as well. These results extend the effects of SETX loss to proteins involved in proteostasis and metabolic homeostasis.

To compare RNA and protein changes, we integrated the proteomic and RNA-seq datasets. Of the 693 DEPs, 144 (approximately 21%) also met the differential expression criteria at the RNA level (**Figure 8C**). Among these, 95 showed concordant changes: 43 were increased and 52 were decreased in both datasets. Concordantly decreased genes included NRXN1, SYN3, MAPT, NTM, ABLIM1, KIF1B, KIF3C, CADPS2, GRIA1, and multiple PCDHA family members (**Figure 8D**). These findings provide evidence at both the RNA and protein levels for altered neuronal structural, adhesion, and synaptic programs. The remaining 49 overlapping genes showed opposite changes in RNA and protein abundance. Twenty showed decreased RNA but increased protein, whereas 29 showed increased RNA but decreased protein (**Figure 8C**). An additional 549 DEPs were not identified as DEGs in the RNA-seq analysis. Thus, only a subset of protein changes was accompanied by statistically significant RNA changes under the thresholds used.

Together, the transcriptomic and proteomic analyses identify shared alterations in neuronal development, axonal and synaptic organization, and cellular homeostasis. They also reveal protein-level changes that are not captured by the significant RNA-level changes, indicating that the two datasets provide complementary views of the effects of SETX loss.

## DISCUSSION

SETX is a multifunctional RNA/DNA helicase involved in transcription termination, R-loop resolution, RNA processing, DNA damage responses, and maintenance of genome stability [4, 46, 47]. Despite these broadly important cellular functions, pathogenic *SETX* mutations preferentially affect the nervous system, causing distinct neurological disorders including ALS4 and AOA2 [14, 48]. How dysfunction of a ubiquitously expressed protein produces pronounced vulnerability in specific neuronal populations remains poorly understood. Here, we examined SETX loss during human MN development using isogenic hiPSC-derived, knockdown, and direct-conversion models. SETX loss slowed proliferation in fibroblasts, hiPSCs, and NPCs, but these cells could be maintained in culture. In contrast, SETX-deficient MNs showed impaired neurite development, reduced electrical activity, and poor survival. RNA hybrid accumulation showed the largest relative increase in MNs among the cell types examined. RNA-seq revealed altered neurodevelopmental gene expression at the NPC stage and during MN differentiation. Proteomic analysis further identified changes in proteins associated with neuronal structure, synaptic function, and cellular homeostasis. Together, these findings support a stage-dependent requirement for SETX in human MN maturation and maintenance (**Figure 9**).

**Figure 9.**
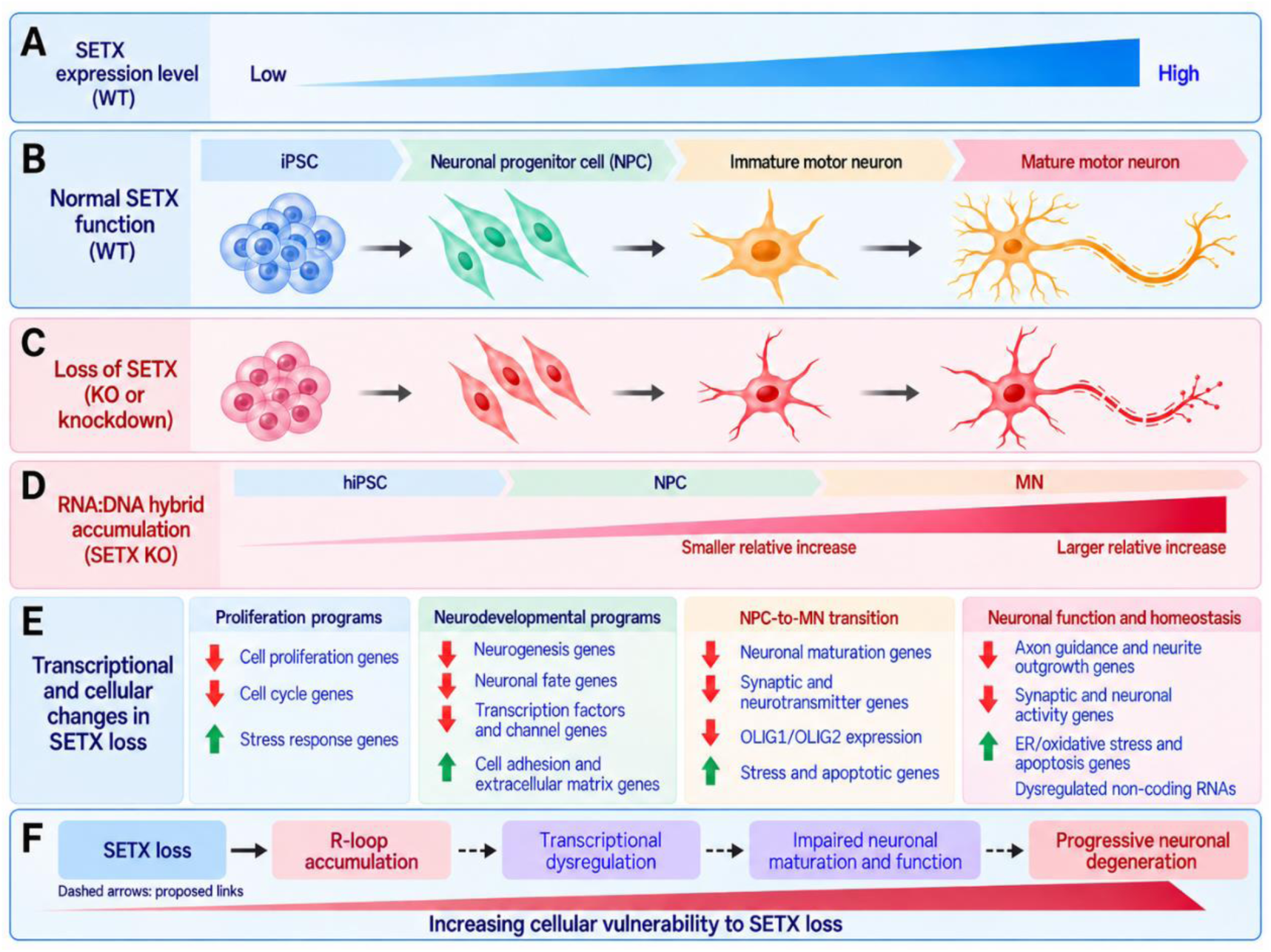
Proposed model of SETX function during human motor neuron differentiation and maturation. (**A**) SETX expression increases during differentiation from human induced pluripotent stem cells (hiPSCs) to neural progenitor cells (NPCs) and motor neurons (MNs). (**B**) Normal MN differentiation from hiPSCs through NPCs and immature MNs to mature MNs. (**C**) SETX knockout or knockdown permits acquisition of MN markers but impairs neurite development and neuronal survival. (**D**) Proposed increase in RNA hybrid accumulation following SETX loss during differentiation. The KO-to-WT increase is greater in MNs than in NPCs. The smaller increase depicted in hiPSCs is hypothesized and was not directly measured. The gradient is schematic and does not represent quantitative values. (**E**) Summary of transcriptional and cellular changes associated with SETX loss. These include reduced proliferation, altered neurodevelopmental gene expression in NPCs, and changes in neuronal maturation, neurite development, synaptic function, and neuronal activity programs during MN differentiation. Stress- and apoptosis-associated genes are increased, while extracellular matrix genes and non-coding RNAs show altered expression. (**F**) Proposed model linking SETX loss to R-loop accumulation, transcriptional dysregulation, impaired neuronal maturation and function, and progressive degeneration. The solid arrow indicates increased R-loop accumulation following SETX loss. Dashed arrows indicate proposed downstream links that remain to be tested. The red gradient illustrates the proposed increase in cellular vulnerability during MN differentiation and maturation. This graphical model was generated with the assistance of ChatGPT (OpenAI) and subsequently reviewed and revised by the authors.

A key finding of this study is that the effects of SETX loss differ between proliferating cells and postmitotic MNs. SETX KO hiPSCs and NPCs proliferated more slowly but could be maintained in culture. KO hiPSCs retained pluripotency-associated features and multilineage differentiation potential, and KO NPCs could generate MNs. However, these MNs developed short, dystrophic neurites and showed progressive neuronal loss. Degeneration was particularly rapid without astrocyte support. These findings suggest that the demands of neuronal differentiation and maturation may increase vulnerability to SETX loss.

Several features of postmitotic neurons may contribute to this vulnerability. Neurons must maintain gene expression and genome stability over long periods without cell division. Their extensive processes require sustained RNA processing, intracellular transport, energy production, and protein quality control [49, 50]. Differences in DNA repair between dividing cells and postmitotic neurons may also influence their response to SETX loss. We found that the KO-to-WT ratio of nuclear S9.6 signal intensity was significantly higher in MNs than in fibroblasts or NPCs, indicating a larger relative increase in RNA hybrid accumulation. Studies in spinal muscular atrophy (SMA) have also linked reduced SETX levels to R-loop accumulation and DNA double-strand breaks [15]. Together, these observations suggest that altered R-loop homeostasis may contribute to the vulnerability of SETX-deficient MNs during differentiation and maturation (**Figures 9D**).

Astrocyte co-culture prolonged the survival of SETX-deficient MNs, showing that the culture environment influences the effects of SETX loss. However, these neurons still showed abnormal neurite morphology and reduced electrical activity. Thus, SETX deficiency appears to create an intrinsic vulnerability. Astrocyte support improved survival but did not prevent structural and functional deficits. The mechanisms underlying this survival benefit remain to be determined.

Our findings distinguish the acquisition of MN markers from subsequent morphological maturation and maintenance. SETX KO cells expressed HB9/MNX1 and ChAT, indicating that SETX loss did not prevent the development of key MN features under our differentiation conditions. However, surviving KO MNs had shorter neurites, dystrophic swellings, and progressive structural deterioration. Thus, MN marker expression alone does not establish normal maturation or long-term neuronal integrity. MEA recordings provide functional evidence supporting this distinction. WT cultures developed spontaneous firing, burst activity, and synchronized network activity. In contrast, SETX KO cultures showed reduced firing rates, lower spike amplitudes, fewer bursts, and almost no synchronized network bursts. Thus, SETX loss compromises not only neuronal morphology but also the acquisition of functional properties characteristic of mature neuronal networks. Importantly, these electrophysiological abnormalities could reflect several interconnected processes. Reduced neurite complexity and impaired synaptic development may limit formation of functional neuronal networks, while transcriptional alterations in ion channels, neurotransmitter transporters, and synaptic proteins may directly impair intrinsic excitability and synaptic transmission. Progressive neuronal stress and degeneration may further exacerbate these abnormalities. Our data therefore does not establish whether electrophysiological dysfunction is a primary consequence of SETX loss or secondary to impaired structural maturation. Rather, they indicate that SETX deficiency disrupts multiple coordinated components of the maturation program required to generate functionally competent human MNs.

A major strength of this study is the similar neuronal defects observed across three complementary models. SETX knockdown using independent shRNAs reproduced the neurite abnormalities and neuronal loss observed in hiPSC-derived KO MNs. Knockdown had little effect during the first several days of differentiation, but rapid degeneration followed. Without astrocyte support, KD MNs survived longer than KO MNs, suggesting that residual SETX expression may delay degeneration. This observation supports a dose-dependent requirement for SETX in maintaining human MN integrity, although direct quantitative relationships between SETX protein abundance and neuronal survival will require further investigation. The direct-conversion experiments provide additional support for a neuron-intrinsic role of SETX. Directly converted MNs derived from SETX KO fibroblasts acquired MN markers with an efficiency comparable to WT cells, yet failed to undergo normal morphological maturation. Instead of developing the characteristic elongated neuronal processes observed in WT diMNs, SETX KO cells exhibited smaller soma, shortened and highly branched neurites, arrested neurite extension, and increased dystrophic swellings during the maturation period. Direct conversion bypasses the pluripotent state and much of the conventional NPC differentiation program [34, 51–54]. Thus, reproduction of the phenotype in this independent system argues against the possibility that the abnormalities observed in hiPSC-derived MNs arise solely from defects acquired during pluripotency or early neural induction. Together, the KO, knockdown, and direct-conversion experiments converge on a common conclusion: SETX is dispensable for the initial acquisition of several MN lineage characteristics under our experimental conditions but becomes increasingly important for subsequent morphological maturation and neuronal maintenance.

Our transcriptomic analyses provide a molecular framework for these cellular defects. Even at the NPC stage, SETX KO cells exhibited extensive transcriptional alterations, including downregulation of genes associated with neurogenesis, neuronal fate commitment, and nervous system development. Thus, the molecular consequences of SETX loss begin before overt neuronal degeneration. Importantly, SETX KO NPCs remained capable of neural differentiation, indicating that these early transcriptional abnormalities alter developmental competence rather than completely blocking neural lineage progression.

The transition from NPCs to MNs revealed a more pronounced divergence between WT and SETX KO cells. During WT differentiation, expression of cell-cycle and DNA-replication genes decreased, while genes associated with neurite growth, synaptic development, and neuronal function increased. SETX KO cells showed a broadly similar transition, but several neuronal and stress-response programs differed. Some genes associated with neuronal function showed weaker induction or met the upregulation criteria only in WT cells. Other developmental and stress-associated genes showed greater induction or met those criteria only in KO cells. These findings suggest that SETX loss alters the coordination of gene expression during MN differentiation. OLIG1 and OLIG2 were among the developmental regulators downregulated during KO differentiation. OLIG2 is a central determinant of spinal MN progenitor identity and participates in the transcriptional network controlling MN specification [16, 55]. The strong reduction of OLIG1/2 during the SETX KO differentiation process raises the possibility that SETX deficiency perturbs transcriptional programs established during early MN specification, with consequences that become apparent during later maturation.

A second major transcriptomic signature of SETX deficiency was activation of cellular stress pathways. SETX KO MNs exhibited increased expression of genes associated with ER stress, oxidative stress, detoxification, apoptosis, and extracellular remodeling. Several of these stress-associated programs were already preferentially activated during the NPC-to-MN transition, indicating that cellular stress emerges as SETX-deficient cells acquire neuronal identity. This observation is consistent with the established biochemical functions of SETX. SETX resolves RNA/DNA hybrids and participates in transcription termination, RNA metabolism, transcription– replication conflict resolution, and genome maintenance [4, 9, 11, 12, 56]. Disruption of these processes can generate transcriptional stress, DNA damage, proteostatic abnormalities, and downstream cellular stress responses. Previous studies have linked SETX deficiency to abnormal R-loop accumulation and DNA damage in several contexts, including SMA, whereas other studies have implicated mitochondrial function, and stress-granule regulation [15, 57–59]. These mechanisms are not mutually exclusive and may converge to compromise neuronal homeostasis.

The disease implications also require a distinction between SETX loss and disease-associated variants. Biallelic SETX mutations cause AOA2, whereas dominant mutations cause ALS4 [1, 13, 14]. Complete knockout defines consequences of SETX deficiency but is not a direct model of ALS4 or of the full clinical phenotype of AOA2. Studies of specific ALS4 variants have reported reduced R-loop abundance and altered regulation of SETX-dependent hybrid resolution [23, 56]. Thus, increased hybrid accumulation after knockout should not be assumed to explain all SETX-associated diseases. Instead, these studies and our findings support the broader possibility that disturbances in SETX-dependent RNA regulation can compromise neuronal function through different molecular routes. Direct comparisons between knockout and disease-associated variants will be needed to identify shared downstream pathways and mutation-specific effects.

## ACKNOWLEDGMENTS

We thank members of the Ding laboratory for their technical assistance, Dr. David Gross for the assistance in dot-blotting assay, and Drs. Armando Salinas, Lucy Robinson, Shile Huang, and Xiuping Yu for helpful discussions. We also acknowledge support from LSU Health Shreveport Core Facilities (RRID:SCR_024775) and the Redox Molecular Signaling Core (RRID:SCR**_**024778) for proteomics analysis.

## AUTHOR CONTRIBUTIONS

B.D. conceptualized and designed experiments. M.A.H., B.D., M.K.I., Y.D., H.C. X.S., T.K. and S.L. performed experiments. B.D. and M.A.H. analyzed and interpreted data. B.D. wrote the original manuscript. B.D., M.A.H. and S.L. edited the manuscript. All authors have read and provided inputs to the manuscript.

## DATA AVAILABILITY

Reserved DOI available on FigShare and will be made public at time of publication at: https://doi.org/10.6084/m9.figshare.34948873

## MATERIALS AND METHODS

### Cell lines and culture conditions

The use of human cell lines, including human induced pluripotent stem cells (hiPSCs), in this study was approved by the Institutional Review Board (IRB) of Louisiana State University Health Sciences Center at Shreveport (LSUHSC-S, IRB ID: STUDY00002945). All cell lines used in this study were listed in **Supplemental Table S1**. HEK 293T cells (CRL-11268) were purchased from ATCC. A SETX KO fibroblast line was generated from an immortalized human fibroblast cell line (R2F/HTERT). Briefly, R2F fibroblasts were transduced with a lentiviral CRISPR/Cas9 vector targeting a 20-nucleotide protospacer sequence (5′-TGTTGAAGCACTTTGTCGGA-3′) within exon3 of the SETX gene. Two weeks after the transduction, single cells were deposited into individual wells of 96-well plates containing conditioned medium (spent media harvested from cell cultures) using FACS. Genomic DNA was isolated from the single-cell clones and sequenced across the Cas9 cutting site of the SETX gene to identify clones with biallelic frameshift indels. The absence of SETX protein expression was further confirmed by Western blot.

SETX KO hiPSC lines and corresponding WT isogenic controls were generated from the healthy hiPSC line WTC11 as previously described [25–28]. All hiPSCs were maintained in complete mTeSR1 medium (STEMCELL Technologies) on Matrigel (Corning) coated dishes at 37 °C and 5% CO_2_ with saturating humanity, and the medium was daily replaced.

The recipes of cell culture medium were:

1. HEK medium: DMEM (Gibco) supplemented with 10% fetal bovine serum (FBS, Corning, NY, USA) and 1% penicillin/streptomycin (ThermoFisher).
2. Fibroblast medium: DMEM supplemented with 15% FBS and 1% penicillin/streptomycin.
3. hiPSC medium: complete mTeSR1 medium and 1% penicillin/streptomycin.
4. Neural progenitor cell (NPC) medium: DMEM/F12 and neurobasal medium (1:1) containing 0.5% N2, 1% B27, 1% GlutaMax, 1% NEAA, 50 µM β-ME, 1% penicillin/streptomycin, 10 ng/mL EGF and 10 ng/mL bFGF.
5. Neuronal maturation medium: DMEM: F12: neurobasal (2:2:1), 1% N2 (Invitrogen), 1% B27 (Invitrogen), 1% penicillin/streptomycin, and supplemented with 5 mM FSK and 10 ng/mL each of BDNF, GDNF, and NT3 (PeproTech).

### Plasmid construction and lentivirus production

A third-generation lentiviral vector (*pCSC-SP-PW-IRES-GFP*) was used to express reprogramming factors NEUROG2-IRES-GFP-T2A-SOX11 and NEUROG2-IRES-SOX11 for direct conversion fibroblast cells into MNs, and NGN2-IRES-ISL1-T2A-LHX3 (Addgene #221814) for generation of iPSC-derived MNs as previously reports [29, 32, 34, 60]. Another third-generation lentiviral vector (*LV-CAG-mCherry-miRE-Luc*) was used to express shRNAs targeting SETX as described in previously reports [34, 60, 61]. Briefly, shRNA oligos targeting the gene coding regions were synthesized, PCR amplified with flanking miR-30 sequences, and ligated into the above lentiviral vector at the *XhoI* and *EcoRI* restriction sites. The targeting sequences (gRNAs) were listed in **Supplemental Table S2.** All plasmids were verified by restriction enzyme digestions and DNA sequencing. Each of these vectors was co-transfected with packaging plasmids (Addgene #12251, #12253, and #12259) into HEK293T cells for virus production [62, 63]. Replication-incompetent lentiviruses were produced, and viral supernatants were collected at 48 hrs and 72 hrs post-transfection as previously described [64]. The viral supernatants were filtered through 0.45 μm syringe filters and stored at 4 °C prior to cell transduction.

### Embryoid bodies (EB) formation

As previously described [25, 34, 65], cultured hiPSCs were dissociated with Versene and transferred to low attachment 10-cm petri dishes in KOSR medium (DMEM/F12 medium containing 20% KnockOut Serum Replacement, 1% GlutaMax, 1% non-essential amino acids, 50 μM β-mercaptoethanol and 1% penicillin-streptomycin) supplemented with 10 µM Y-27632. The medium was changed every other day and EBs gradually formed. After 7 days of suspension culture, EBs were digested with 0.25% Trypsin and cultured on gelatin-coated plates with KOSR medium for another 7 days. The total RNAs were extracted for RT-PCR analysis.

### Direct conversion of MNs from human fibroblasts

Direct conversion of fibroblasts into MNs using lentiviral delivery of 4 factors (NGN2, SOX11, ISL1 and LHX3) as previously described [32, 34].

### Generation of hiPSC-derived MNs

MNs were prepared from hiPSCs as previously described [29, 30, 66]. Briefly, hiPSCs were cultured in mTeSR1 medium with 10 µM all-trans-retinoic acid (RA, Sigma) and 0.5 mM VPA in Matrigel-coated 6-well plates for 7 days. Cells were then digested with Versene and gently pipetted into small clumps supplemented with 10 μM Y-27632 (STEMCELL Technologies). Cell clumps were cultured in KOSR medium for 4 days, followed by cultured in NSP medium for another week. The neurospheres were then dissociated into single cells with Accutase (Innovative Cell Technologies) and maintained in neural progenitor cell medium. For MN differentiation [26, 29, 67], neural progenitor cells were plated into Matrigel-coated plates at a density of 3×10^4^ cells/cm^2^ and transduced with a lentiviruse expressing *NEUROG2-IRES-ISL1-T2A-LHX3*. Culture medium was replaced the next day with neuronal maturation medium. Neurons were dissociated with Accutase on day 5 and replated onto Matrigel-coated coverslips with or without the presentence of astrocytes depending on desired experiments. The medium was half changed twice a week until analysis.

### Immunocytochemistry (ICC)

ICC analysis was performed as previously described [34, 60, 68]. Briefly, Cultured cells at the desired time points were fixed with 4% paraformaldehyde (PFA) in PBS for 15 min at room temperature (RT) and then permeabilized and blocked for 1 hr in blocking buffer (PBS containing 0.2% Triton X-100 and 3% BSA). They were subsequently incubated overnight with primary antibodies in blocking buffer at 4 °C, followed by washing and incubation with corresponding fluorophore-conjugated secondary antibodies. Cell nuclei were counterstained with Hoechst 33342 (HST, ThermoFisher). Primary antibodies used in this study were listed in **Supplemental Table S3**.

### Imaging and quantification analysis

Images were obtained with Nikon A1R, Leica TCS SP5, Zeiss LSM700 (63×/1.4 NA) confocal microscopes or an inverted fluorescence microscope (Olympus, CKX53). For quantification of fluorescence intensity of transport reporters, as large an area as possible was measured within the nucleus or cytoplasm, as previously described [33, 69]. An unbiased approach for data collection was employed. The person who analyzed the images was completely blinded to the sample information. The “mean” values were used to quantify an average signal intensity.

### MEA recording and data analysis

Spontaneous activity of cultured MNs was recorded using a multi-electrode array (MEA) system. MEA plates (24-well CytoView MEA plates with 16 PEDOT electrodes per well, Axion BioSystems) were pre-coated with Matrigel to support neuronal adhesion. iPSC-derived MNs were seeded directly onto the recording electrode array at a density of 80,000 cells per well, and the wells were supplied with neuronal maturation medium ∼1 hour after seeding. To maximize survival and network formation, MNs were co-cultured with primary astrocytes added directly onto MNs. Wells with comparable cell numbers were verified by bright-field microscopy on each recording day.

Extracellular field potentials were recorded on an Axion Maestro Edge with AxIS Navigator^TM^ software v3.12.7, with the recording chamber maintained at 37 °C and 5% CO_2_. Plates were equilibrated on the recording stage for 5 minutes prior to acquisition to allow stabilization of activity. Recordings of spontaneous activity were collected at multiple time points across the maturation window of 7 DPI start day through 42 DPI end day. The same wells were tracked longitudinally for each genotype.

Spike detection was performed using the adaptive threshold-crossing algorithm in AxIS Navigator^TM^ software v3.12.7 with a detection threshold of 6× SD of the rolling baseline noise. Active electrodes were defined as those exhibiting ≥ [5 spikes per minute]. Single-electrode bursts were identified using the inter-spike interval (ISI) threshold method with parameters of a minimum of 5 spikes per burst and a maximum ISI of 100 ms. Network bursts (synchronous activity across multiple electrodes within a well) were detected using the Inter-burst Interval algorithm with thresholds of at least 35% of active electrodes participating, a minimum of 50 spikes per network burst, and a maximum of 100 ms ISI. All spike sorting and burst detection parameters were applied identically to SETX-KO and isogenic control wells.

Further recordings were analyzed by Neural Metric Tool, v4.1.5. Afterwards, the following parameters were extracted on a per-well basis: mean firing rate (spikes per second); mean spike amplitude (µV); single-electrode burst frequency (bursts per minute); mean burst duration (seconds); network burst number per recording; and network burst frequency (network bursts per minute). Continuous waveform plots and raster plots were generated for representative wells to visualize spike activity and network synchrony, respectively.

### R-loop assay by Dot blot

RNA-DNA hybrids (R-loops) were detected using an S9.6 antibody-based dot blot assay [37, 38]. Briefly, genomic DNA was extracted from cultured cells using the Quick-DNA™ Miniprep Kit (Zymo). DNA concentrations were determined using a NanoDrop. Aliquots were treated with RNase H served as negative controls. Genomic DNA with a serial dilution was manually spotted onto positively charged nylon membranes and air-dried at room temperature. Following UV crosslinking, membranes were blocked with 5% bovine serum albumin (BSA) in Tris-buffered saline and sequentially incubated with the S9.6 monoclonal antibody (1:1,000) and an appropriate horseradish peroxidase (HRP)-conjugated secondary antibody. Immunoreactive signals were detected using enhanced chemiluminescence (ECL) and imaged using a Bio-Rad ChemiDoc imaging system. A separate membrane containing matched genomic DNA samples was stained with 0.02% (w/v) methylene blue in 0.3 M sodium acetate buffer (pH 5.2) served as DNA loading. Dot intensities were quantified using ImageJ software following background subtraction, and S9.6 signals were normalized to the corresponding methylene blue staining intensities.

### RNA isolation, sequencing, and Bioinformatics Analysis

SETX-knockout (KO) and isogenic control hiPSC-derived motor neurons (MNs) were generated as described above and cultured in MN maturation medium until 4 days post-induction (DPI). Three independent biological replicates were collected for each genotype (n = 3 SETX-KO; n =3 isogenic control). Cultures were rinsed once with ice-cold phosphate-buffered saline (PBS), and total RNA was isolated using the PureLink^TM^ RNA Mini Kit (Invitrogen) according to the manufacturer’s instructions. RNA concentration and purity were measured on a NanoDrop instrument. The RNA integrity was determined by the Agilent 2100 BioAnalyzer (Agilent Technologies) and sequenced using the Illumina NovaSeq (Novogene). The data obtained in FASTQ files from RNA sequencing were aligned to the Ensembl hg38 human genome using the HISAT2 program. Gene-level abundances were estimated as FPKMs (Fragments Per Kilobase of transcript sequence per Million base pairs sequenced). The read count summarization to each gene (i.e., counts) was calculated using featureCounts. Genes with fewer than 10 (low-count threshold) total reads across all samples were filtered out before analysis. Analyses of differential expressions of transcripts were performed with DESeq2 v1.48.2 (Genes with a false discovery rate (FDR) value of <1% and an adjusted *P* value < 0.05 and an absolute fold change ≥1.5 (|log2FC| ≥ 0.585) were considered to be differentially expressed genes (DEGs). Volcano plots and heat maps were created using the Rgplot, ggplot2, and pheatmap / ComplexHeatmap packages in RStudio 2025.09.1 Build 401.

Gene Ontology (GO) enrichment analysis was performed using the clusterProfiler R package (version 4.16.0) [70]. Gene symbols were converted to Entrez Gene IDs using the org.Hs.eg.db annotation package. Enrichment analyses were conducted separately for GO Biological Process (BP), Molecular Function (MF), and Cellular Component (CC) categories using the enrichGO function. The background consisted of all genes with GO annotations available in org.Hs.eg.db. Statistical significance was assessed using the Benjamini-Hochberg method for multiple-testing correction, and GO terms with a false discovery rate (FDR)-adjusted *P* value < 0.05 were considered significantly enriched.

Protein-protein interaction (PPI) networks were constructed using the STRING database to investigate functional associations among proteins encoded by DEGs [71]. Gene symbols were mapped to their corresponding proteins in the human STRING database (*Homo sapiens*, taxonomic ID: 9606). High-confidence protein associations were selected using a minimum combined interaction score of 0.700. Network topology was analyzed in R using the igraph package [72]. Network characteristics, including the number of connected proteins, interactions, connected components, network density, and node degree, were calculated. Node degree was defined as the number of interactions associated with each protein. Network communities were identified using the Louvain community detection algorithm, and the seven largest modules were highlighted for visualization. Networks were visualized using the ggraph and ggplot2 packages, with node size proportional to degree and node color representing module membership. Smaller network modules were displayed in gray. Major modules were assigned descriptive biological labels based on their constituent genes and associated biological functions.

### Protein extraction and western blotting analysis

Phenol extraction method was used for protein extraction to minimize protein degradation. Briefly, cells were harvested and cell pellets were washed once with ice-cold phosphate-buffered saline (PBS). The washed cell pellets were resuspended in 100 µL of ice-cold PBS, followed by the addition of 5 µL of β-mercaptoethanol and 100 µL of molecular biology-grade phenol equilibrated to pH 8.0. The samples were vortexed for 30 s to lysis cells. Proteins were subsequently precipitated by adding 1.2 mL of methanol containing 0.1 M ammonium acetate, followed by thorough mixing. The samples were centrifuged at 20,000 × g for 30 min at 4°C, and the supernatants were carefully discarded. The resulting protein pellets were washed with 1 mL of methanol containing 0.1 M ammonium acetate by gentle inversion and centrifuged at 20,000 × g for 10 min at 4°C. The protein pellets were subsequently washed with 1 mL of acetone by gentle inversion and centrifuged at 20,000 × g for 10 min at 4°C. Following removal of the acetone, an additional centrifugation at 20,000 × g for 2 min at 4°C was performed to eliminate residual solvent. The protein pellets were air-dried at room temperature for 30 min and dissolved in 100 µL of freshly prepared 1× SDS-PAGE loading buffer containing Tris-HCl (pH 6.8), sodium dodecyl sulfate (SDS), glycerol, bromophenol blue, and β-mercaptoethanol. Then, samples were incubated at 50°C for 30 min instead of boiling. The extracted proteins were subsequently subjected to SDS-PAGE and Western blot analysis or stored at −20°C until further use.

Equal amounts of cell lysates (20 µg per lane) were used for SDS-PAGE and western blot analysis as previously described [34, 68, 73, 74]. Primary antibodies and dilutions were listed in **Supplemental Table S3.** HRP-conjugated secondary antibodies and Clarity Western ECL substrate (Bio-Rad) were used to visualize the protein bands with a BioRad ChemiDoc.

### Mass Spectrometry assay

Cultured neurons were harvested and proteins were extracted using phenol extraction method as described above. Three replicates were prepared in each condition. For LC-MS/MS analysis of these dissolved proteins, 10 µg of protein from each sample was loaded into 12% SDS-PAGE, and then stained with Imperial Protein Stain (ThermoFisher) as previously described [68]. The Gel lanes of each sample were cut and digested according to the manufacturer’s protocol of the Thermo Scientific™ Gel Tryptic Digestion kit. The LC-MS/MS analysis was carried out using an Ultimate 3000 RSLCnano system connected to an Orbitrap Exploris 480 mass spectrometer. Raw data files were processed in Proteome Discoverer v3.3 (Thermo Fisher Scientific), with peptide identification performed using the CHIMISTRY search engine against a species-specific FASTA database (Human Sapiens, version 2026-01-28, taxonomy ID 9606) and the inferys_5.8.0_fragmentation prediction model. The significantly expressed proteins between SETX KO and WT were defined by a threshold of 1.5-fold changes and p-value < 0.05.

### Quantitative real-time PCR analysis

As described previously [60, 68, 75], total RNA was extracted from cultured neurons using TRIzol (Life Technologies). cDNA synthesis reactions were performed using 0.5 µg of total RNA from each sample with the SuperScriptIII First-Strand kit (Life Technologies) and random hexamer primers. Real-time PCR was performed in triplicate using primers, SYBR GreenER SuperMix (Invitrogen), and the BIO-RAD CFX-96 Fast Real-Time PCR system. Target mRNA levels were normalized to the reference gene *GAPDH* or *TUBB3* by 2^−ΔΔCt^ method. Sequences of RT-PCR primers were listed in **Supplemental Table S2.**

### Statistical analysis

Statistical analysis was done in GraphPad Prism. The D’Agostino & Pearson omnibus normality test was conducted first to determine if the data are normally distributed. If the data passed the normality test, one-way or two-way ANOVA was used to determine significance. If the data did not pass the normality test, the Kruskal-Wallis test was used to determine significance. Results are expressed as mean ± SD of at least three biological replicates or three independent experiments, and *P* < 0.05 is considered significant.

