## Supplemental information for "Stage-Dependent Requirement for SETX in Human Motor Neuron Maturation and Survival"

**This PDF file includes:**

Figures S1-S8

Tables S1-S3

Note: Tables S4-S9 will be uploaded as Excel files separately.

### Supplementary Figures and Figure Legends

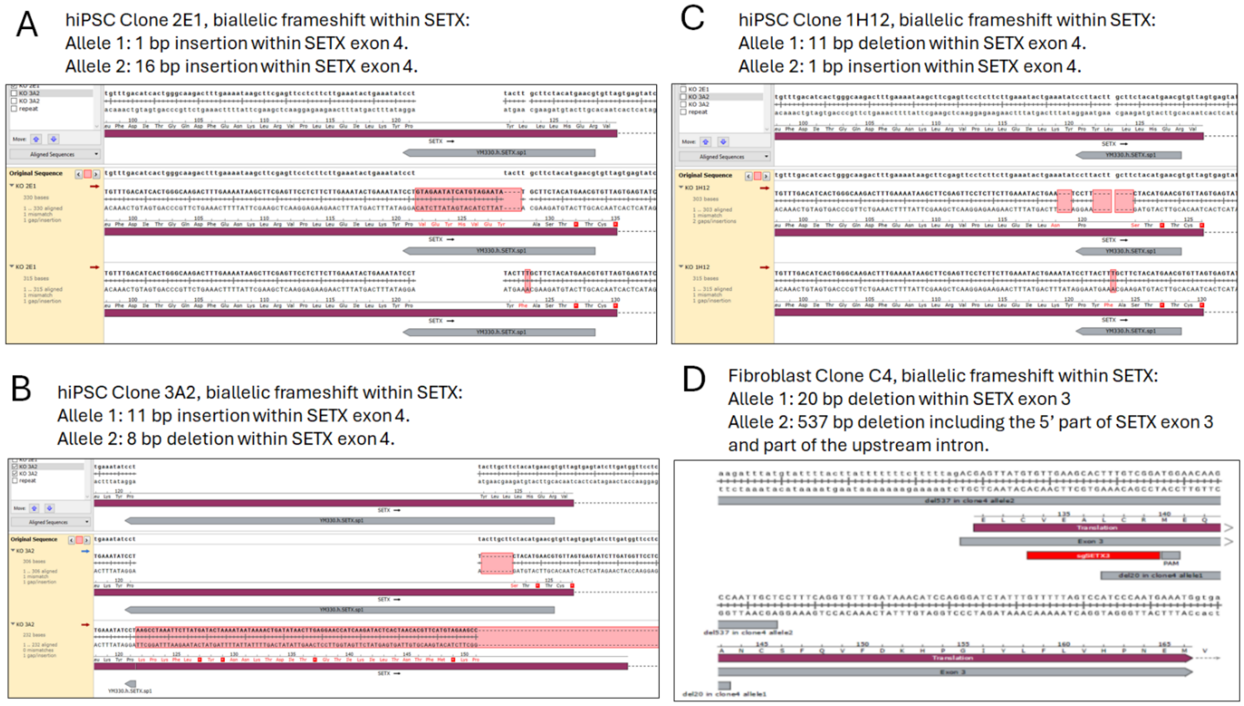


**Figure S1.** DNA sequence and alignment confirmed SETX gene was knocked out with biallelic frameshift in three human induced pluripotent stem cell clones (A-C) and a fibroblast clone (D).

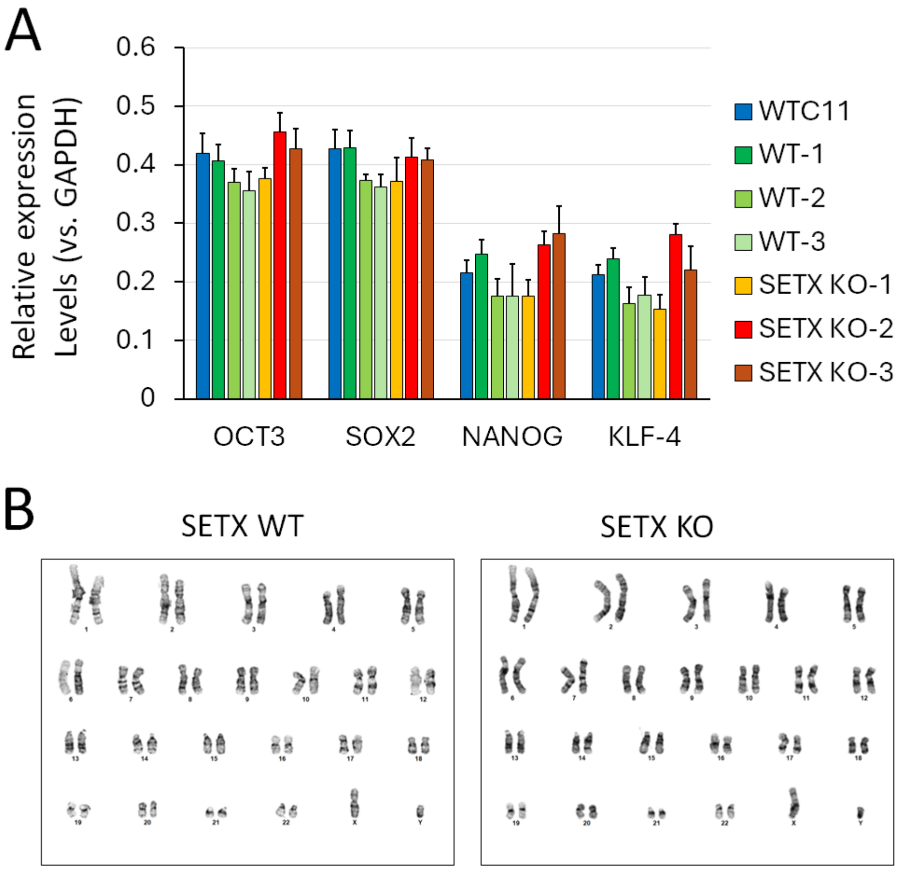


C

| STR | Parental (WTC11) | WT-1 1A3 | WT-2 1A5 | WT-3 1A10 | SETX KO-1 2E1 | SETX KO-2 1H2 | SETX KO-3 3A2 |
|---|---|---|---|---|---|---|---|
| D19S433 | 15, 16 | 15, 16 | 15, 16 | 15, 16 | 15, 16 | 15, 16 | 15, 16 |
| D2S1338 | 17, 18 | 17, 18 | 17, 18 | 17, 18 | 17, 18 | 17, 18 | 17, 18 |
| FGA | 22, 25 | 22, 25 | 22, 25 | 22, 25 | 22, 25 | 22, 25 | 22, 25 |
| PentaE | 13, 15 | 13, 15 | 13, 15 | 13, 15 | 13, 15 | 13, 15 | 13, 15 |
| VWA | 17, 19 | 17, 19 | 17, 19 | 17, 19 | 17, 19 | 17, 19 | 17, 19 |
| CSF1PO | 10, 12 | 10, 12 | 10, 12 | 10, 12 | 10, 12 | 10, 12 | 10, 12 |
| D13S317 | 8, 11 | 8, 11 | 8, 11 | 8, 11 | 8, 11 | 8, 11 | 8, 11 |
| D3S1358 | 15, 16 | 15, 16 | 15, 16 | 15, 16 | 15, 16 | 15, 16 | 15, 16 |
| D8S1179 | 11, 13 | 11, 13 | 11, 13 | 11, 13 | 11, 13 | 11, 13 | 11, 13 |
| TH01 | 6, 7 | 6, 7 | 6, 7 | 6, 7 | 6, 7 | 6, 7 | 6, 7 |
| D16S539 | 9, 10 | 9, 10 | 9, 10 | 9, 10 | 9, 10 | 9, 10 | 9, 10 |
| D21S11 | 29 | 29 | 29 | 29 | 29 | 29 | 29 |
| D5S818 | 9, 10 | 9, 10 | 9, 10 | 9, 10 | 9, 10 | 9, 10 | 9, 10 |
| D7S820 | 10, 12 | 10, 12 | 10, 12 | 10, 12 | 10, 12 | 10, 12 | 10, 12 |
| TPOX | 8, 11 | 8, 11 | 8, 11 | 8, 11 | 8, 11 | 8, 11 | 8, 11 |

**Figure S2. Pluripotency and cell identity verification.**

**(A)** RT-PCR results showing the comparable expression levels of pluripotent markers in newly generated iPSC lines compared with the parental line WTC11. N = three experimental experiments.

**(B)** Representative Karyotype of WT and SETX KO iPSC lines.

**(C)** Short tandem repeat (STR) analysis confirmed the same cell identify of newly generated iPSC lines as the parental line WTC11.

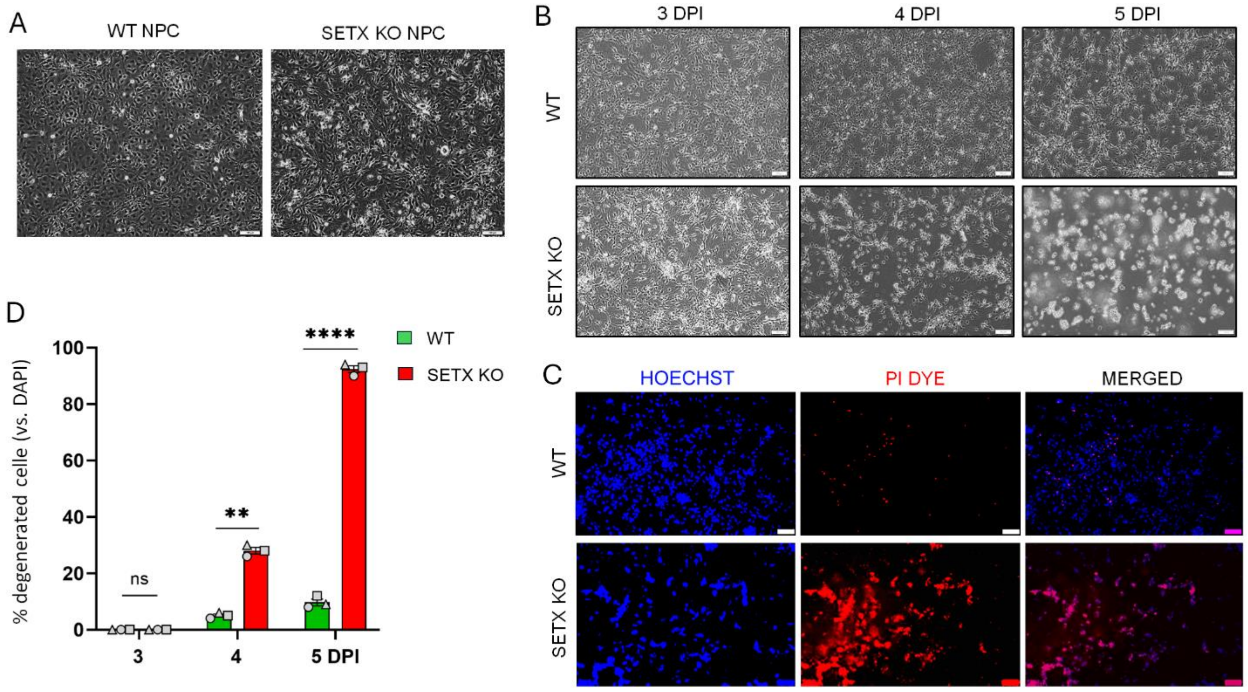


**Figure S3: Cultured SETX KO MNs alone showed premature degeneration.**
(**A**) Micrographs show neuron progenitor cells (NPCs) differentiated from SETX WT (left) and SETX KO (right) iPSCs. Scale bar: 100 µm.
(**B**) Micrographs show differentiated MNs at 3, 4, and 5 DPI (Days post viral infection) induced from WT and SETX KO NPCs. Scale bar: 100 µm.
(**C**) Fluorescent micrographs show Propidium Iodide (PI)-labeled SETX WT and SETX KO MNs at 5 DPI. Hoechst 33342 stained nuclei. Scale bar: 100 µm.
(**D**) Quantification of percentage degenerated MNs at 3, 4, and 5 DPI normalized with DAPI. N (objects) = 3. n (Neurons) > 1000 for each object highlighted with distinct colors. Ns, not significant, ** p<0.01, ****p < 0.0001. Student's t-test.

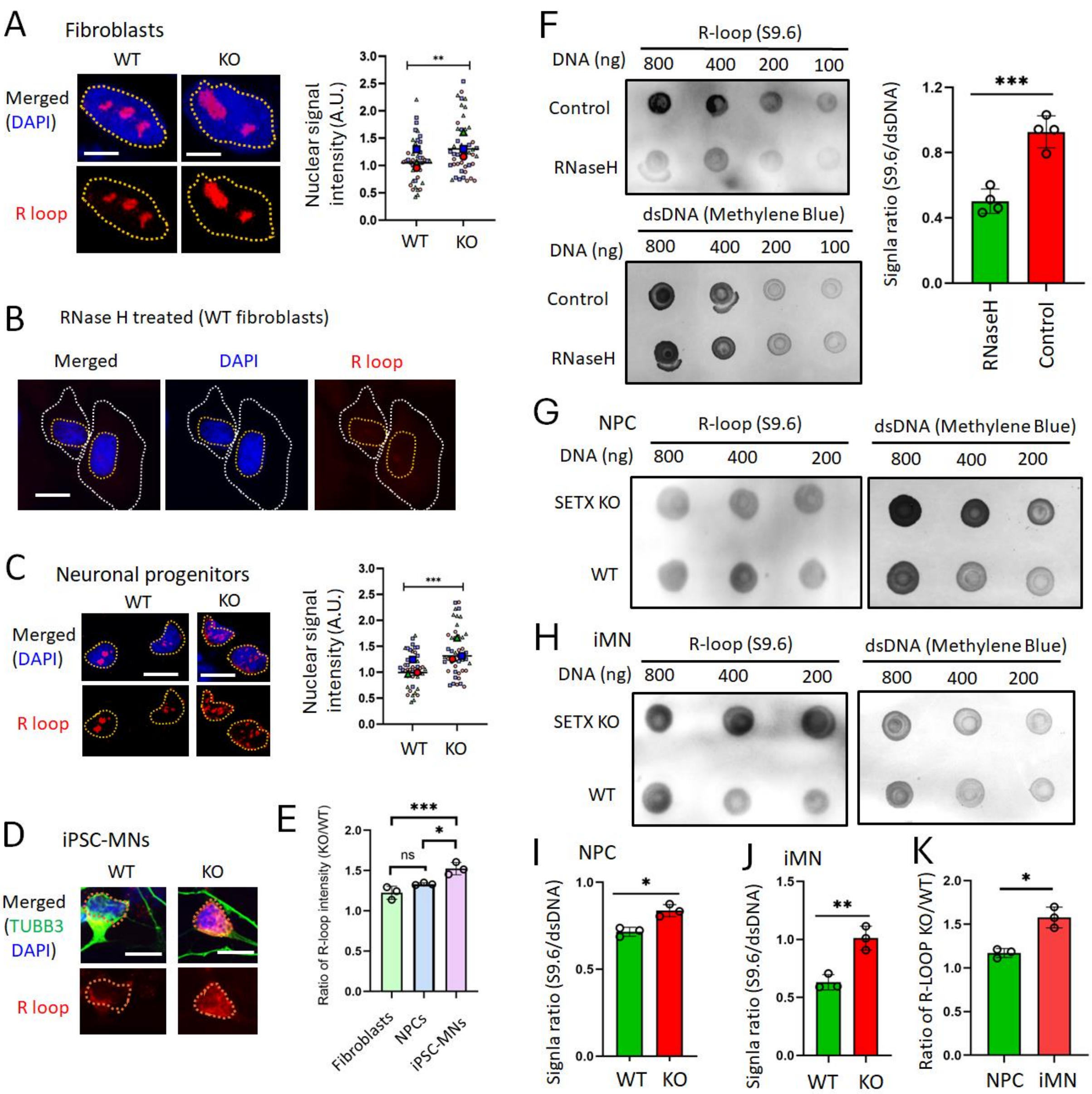


**Figure S4. SETX loss increases RNA hybrid accumulation with the largest relative increase in human MNs.**

(**A**) Representative S9.6 immunofluorescence images and quantification of nuclear signal intensity in WT and SETX KO fibroblasts. Scale bars, 10 µm. n > 50 cells from three independent experiments. **P < 0.01; Student's *t*-test.

(**B**) Representative S9.6 immunofluorescence images of WT fibroblasts following RNase H treatment. Scale bars, 10 µm.

(**C**) Representative S9.6 immunofluorescence images and quantification of nuclear signal intensity in WT and SETX KO NPCs. Scale bars, 20 µm. n > 50 cells from three independent experiments. ***P < 0.001; Student's *t*-test.

(**D**) Representative images of WT and SETX KO hiPSC-derived MNs co-stained for TUBB3 and RNA hybrids using the S9.6 antibody at 4 DPI. Scale bars, 10 µm.

(**E**) KO-to-WT ratios of nuclear S9.6 signal intensity in fibroblasts, NPCs, and MNs. n = 3 independent experiments. ns, not significant; *P < 0.05; ***P < 0.001. One-way ANOVA followed by Dunnett's multiple-comparisons test.
(**F**) S9.6 dot blots of nuclear extracts from HEK cells with or without RNase treatment and quantification of hybrid signal normalized to methylene blue staining of DNA. ***P < 0.001; Student's *t*-test.
(**G, H**) Representative S9.6 dot blots of nuclear extracts from WT and SETX KO NPCs (G) and MNs (H). Methylene blue staining of DNA served as a loading control.
(**I, J**) Quantification of S9.6 dot-blot signals in NPCs (I) and MNs (J), normalized to DNA loading. n = 3 independent experiments. *P < 0.05; **P < 0.01. Student's *t*-test.
(**K**) KO-to-WT ratios of normalized S9.6 dot-blot signal intensity in NPCs and MNs. n = 3 independent experiments. *P < 0.05; Student's *t*-test.

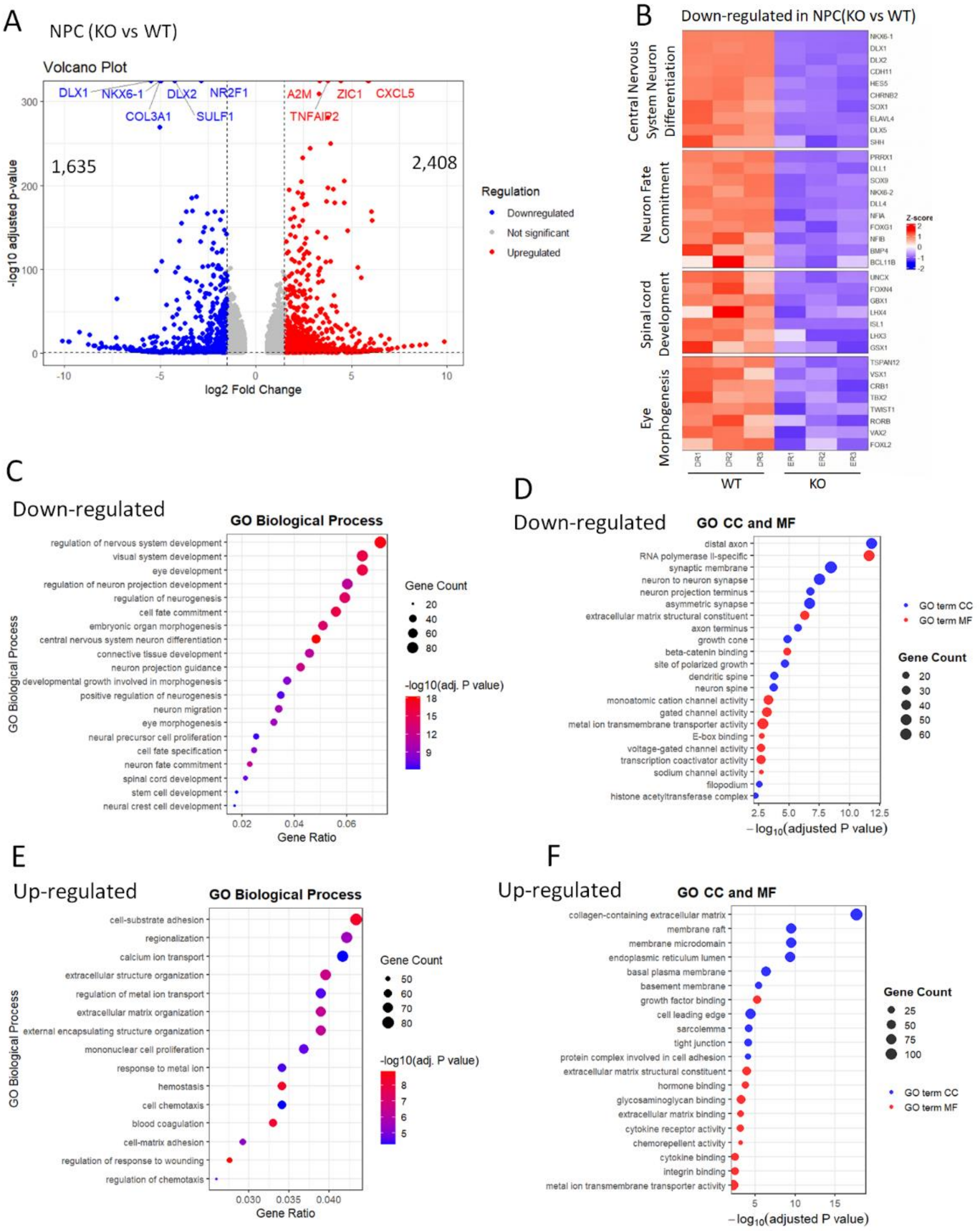


**Figure S5. Transcriptional alterations in SETX KO neuronal progenitor cells (NPC).**
(**A**) Volcano plot showing differentially expressed genes (DEGs) in SETX KO NPCs relative to WT NPCs. Differential expression analysis was performed using DESeq2, with WT NPCs as the reference. Genes with an adjusted *P* value < 0.05 and an absolute fold change ≥1.5 (|log2FC| ≥ 0.585) were considered differentially expressed.
(**B**) Heatmap of selected significantly downregulated genes associated with neuronal development and neurodevelopmental processes.
(**C, D**) GO enrichment analysis of significantly downregulated genes enriched in GO Biological Process (C), and Cellular Component (CC) and Molecular Function (MF) terms (D).
(**E, F**) GO enrichment analyses of significantly upregulated genes enriched in GO BP terms (E), and CC and MF terms (F).

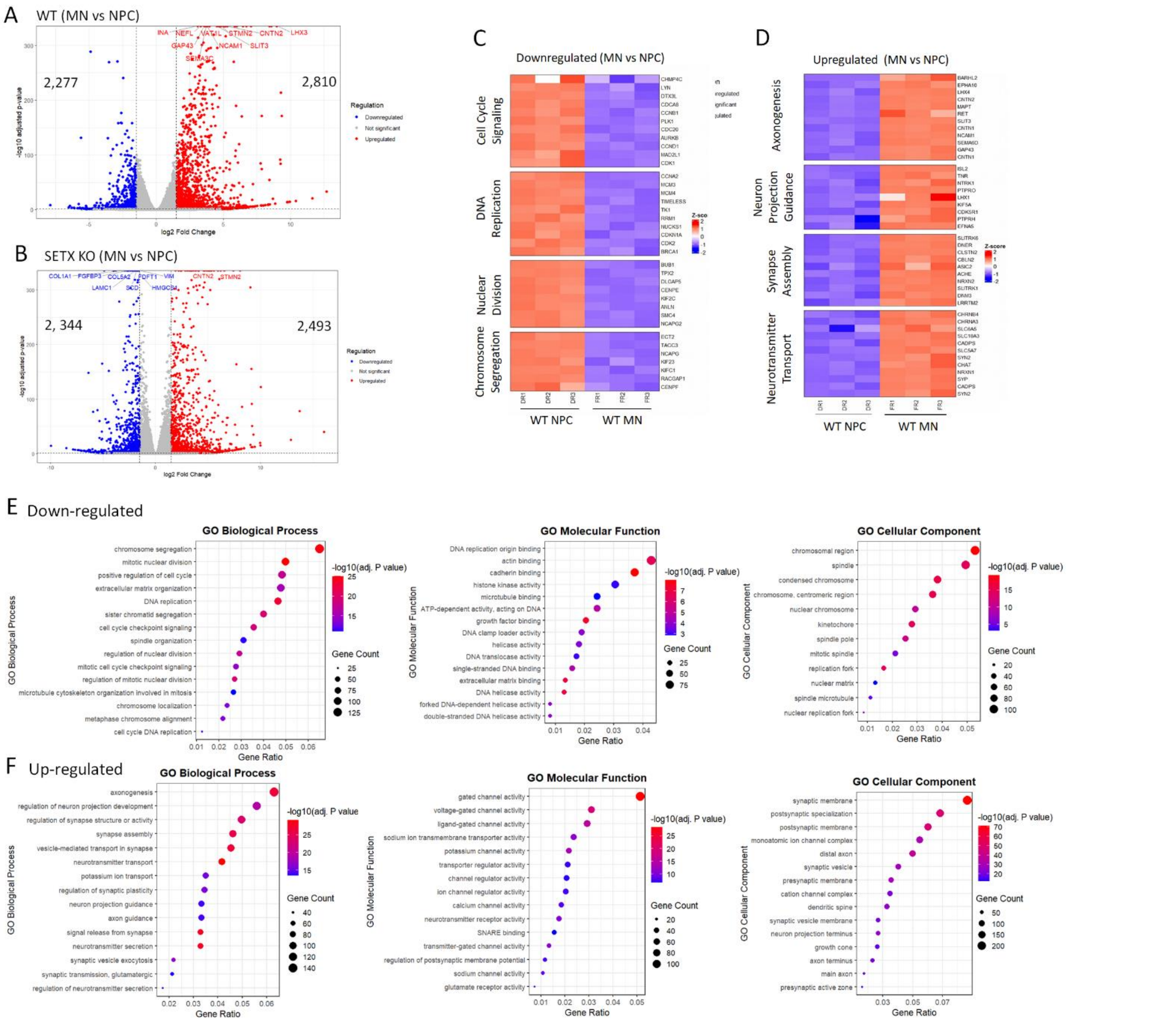


**Figure S6. Transcriptional transition from proliferative NPCs to postmitotic MNs.**
(**A, B**) Volcano plots showing differentially expressed genes (DEGs) in MNs relative to NPCs in WT (A) and SETX KO (B), respectively.
(**C, D**) Heatmaps of selected genes significantly downregulated (C) or upregulated (D) during the NPC-to-MN transition in WT. Genes were manually selected based on biological relevance from significant WT MN-versus-NPC DEGs and organized into functional categories.
(**E, F**) GO enrichment analysis of genes significantly downregulated (E) and upregulated (F) during the NPC-to-MN transition in WT, shown separately for Biological Process (BP), Molecular Function (MF), and Cellular Component (CC) categories.

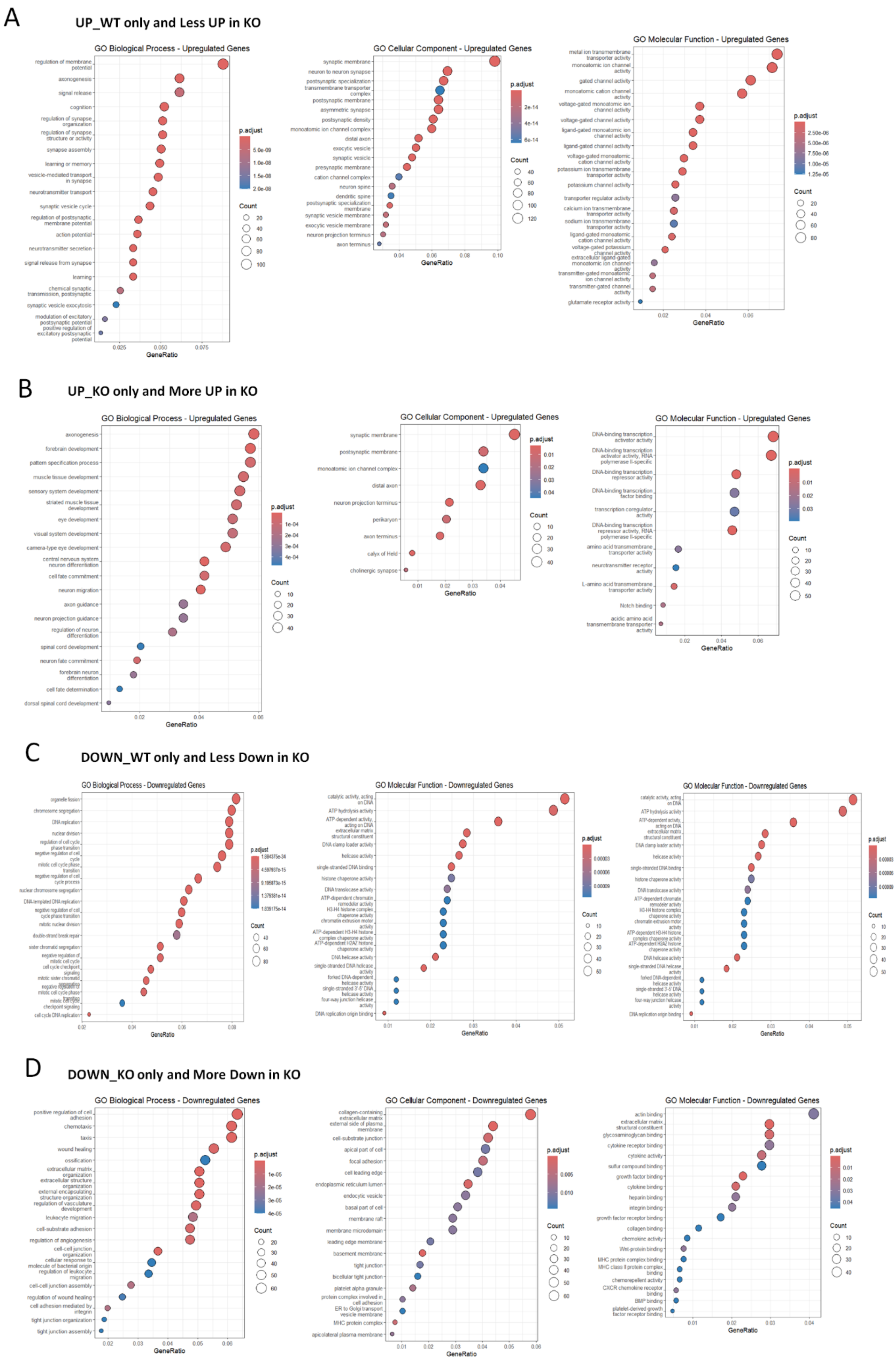

A
UP_WT only and Less UP in KO
GO Biological Process - Upregulated Genes
GO Cellular Component - Upregulated Genes
GO Molecular Function - Upregulated Genes
B
UP_KO only and More UP in KO
GO Biological Process - Upregulated Genes
GO Cellular Component - Upregulated Genes
GO Molecular Function - Upregulated Genes
C
DOWN_WT only and Less Down in KO
GO Biological Process - Downregulated Genes
GO Molecular Function - Downregulated Genes
GO Molecular Function - Downregulated Genes
D
DOWN_KO only and More Down in KO
GO Biological Process - Downregulated Genes
GO Cellular Component - Downregulated Genes
GO Molecular Function - Downregulated Genes

**Figure S7. GO enrichment analysis of genes differentially regulated during the NPC-to-MN transition in WT and SETX KO cells.**

**(A)** GO enrichment analysis of genes that were significantly upregulated during the NPC-to-MN transition only in WT cells or showed a smaller increase in SETX KO cells compared with WT cells.

**(B)** GO enrichment analysis of genes that were significantly upregulated during the NPC-to-MN transition only in SETX KO cells or showed a greater increase in SETX KO cells compared with WT cells.

**(C)** GO enrichment analysis of genes that were significantly downregulated during the NPC-to-MN transition only in WT cells or showed a smaller decrease in SETX KO cells compared with WT cells.

**(D)** GO enrichment analysis of genes that were significantly downregulated during the NPC-to-MN transition only in SETX KO cells or showed a greater decrease in SETX KO cells compared with WT cells.

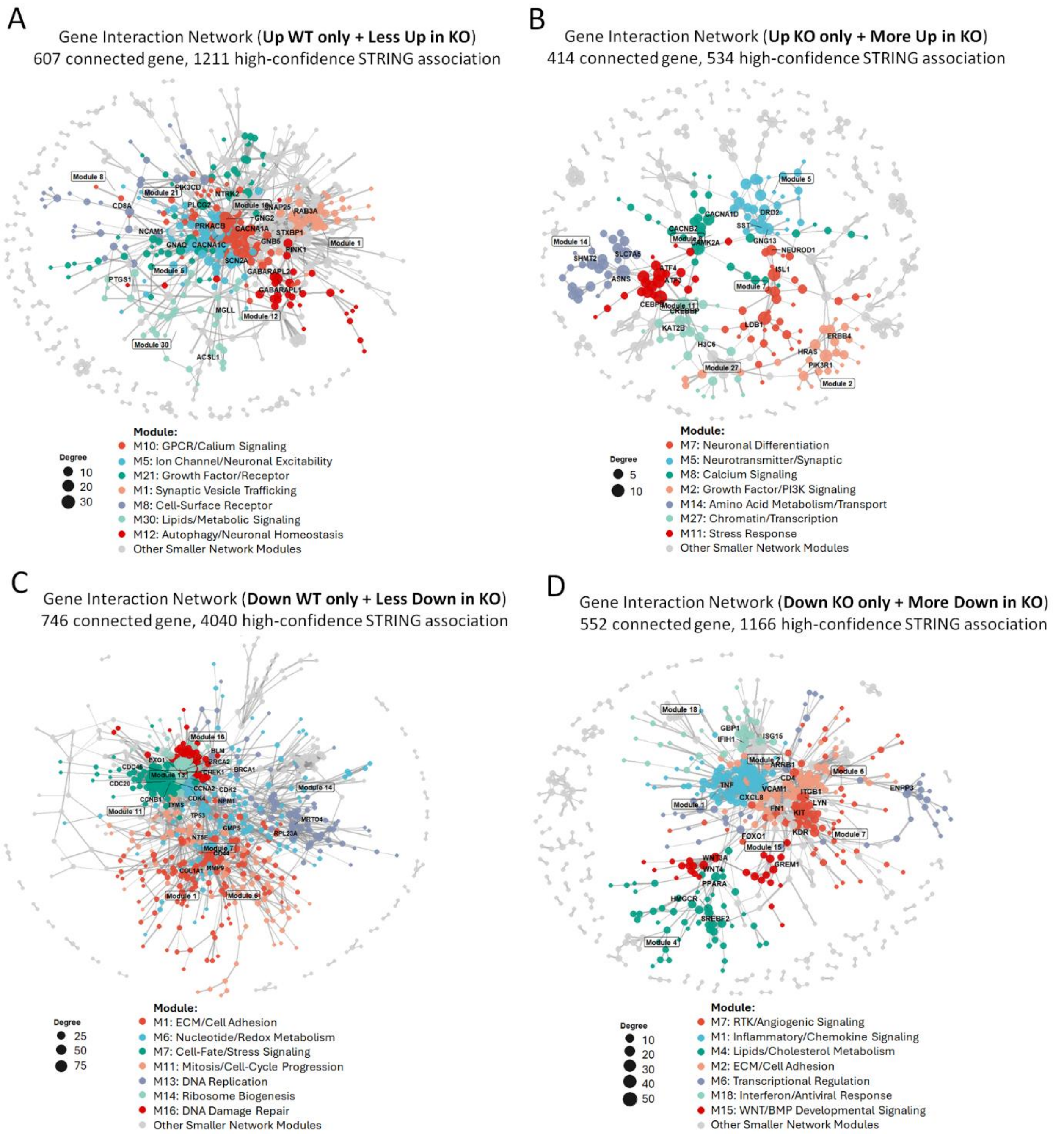


**Figure S8. Gene interaction networks of DEG subgroups differentially regulated during the NPC-to-MN transition in WT and SETX KO cells.**

**(A)** Gene interaction network of genes significantly upregulated during the NPC-to-MN transition only in WT cells or showing a smaller increase in SETX KO cells compared with WT cells.

**(B)** Gene interaction network of genes significantly upregulated during the NPC-to-MN transition only in SETX KO cells or showing a greater increase in SETX KO cells compared with WT cells.

**(C)** Gene interaction network of genes significantly downregulated during the NPC-to-MN transition only in WT cells or showing a smaller decrease in SETX KO cells compared with WT cells.
**(D)** Gene interaction network of genes significantly downregulated during the NPC-to-MN transition only in SETX KO cells or showing a greater decrease in SETX KO cells compared with WT cells.

### Supplementary Tables

**Table S1. List of cell lines used in this study and their Research Resource Identifiers (RRIDs).**

| | Resource | Sex | Age (YR) | Gene Mutations | Note |
|---|---|---|---|---|---|
| **hiPSC lines** | | | | | |
| WTC11 | Coriell Cat# GM25256, RRID:CVCL_Y803 | M | 30 | - | Parental lines |
| WT-1 | This study | M | 30 | - | Clone ID: 1A3 |
| WT-2 | This study | M | 30 | - | Clone ID: 1A5 |
| WT-3 | This study | M | 30 | - | Clone ID: 1A10 |
| SETX KO-1 | This study | M | 30 | Biallelic frameshift | Clone ID: 2E1 |
| SETX KO-2 | This study | M | 30 | Biallelic frameshift | Clone ID: 1H2 |
| SETX KO-3 | This study | M | 30 | Biallelic frameshift | Clone ID: 3A2 |
| **Human fibroblast** | | | | | |
| WT | R2F/HTERT, RRID:CVCL_UZ36 | M | <1 month | Telomerase immortalized cell line | |
| SETX KO | This study | M | <1 month | Biallelic frameshift | |
| **Other cell lines** | | | | | |
| HEK 293T cells | ATCC Cat# CRL-11268, RRID:CVCL_1926 | | | | |

Note: RRIDs, Research Resource Identifiers (RRIDs); M, Male; F, Female.

**Table S2. List of primary antibodies used this study.**

| Name | Source | Catalogue | Host | Immunostaining | Western Blot |
|---|---|---|---|---|---|
| β-Actin | Sigma | A5441 | Ms | - | 1:5000 |
| GAPDH | Proteintech | 50-553-944 | MS | - | 1:2000 |
| GFP | Aves | GFP-1020 | Ck | 1:1000 | 1:5000 |
| MAP2 | Abcam | Ab5392 | Ck | 1:10000 | - |
| NANOG | Abcam | ab21624 | Rb | 1:100 | - |
| OCT4 | Santa Cruz | sc-5279 | Ms | 1:200 | - |
| SETX | Bethyl | A700-050 | Rb | - | 1:300 |
| SOX2 | Santa Cruz | sc-365823 | Ms | 1:200 | - |
| SSEA4 | Abcam | ab16287 | Ms | 1:200 | - |
| TUBB3 | Covance | MMS-435P | Ms | 1:2000 | - |
| TUBB3 | Covance | PRB-435P | Rb | 1:2000 | - |
| γH2Ax | Bethyl | A300081A | Rb | 1:500 | - |
| S9.6 | Millipore-Sigma | 4332229 | MS | 1:300 | - |

Note: Ck, chicken; Gt, goat; Ms, mouse; Rb, rabbit; Sh, sheep; - not determined in this study.

**Table S3. List of Oligonucleotide sequences**

| Name | Sequence (5'-3') | Note |
|---|---|---|
| SETX-shRNA-1 | AAGGACAGAATAGAAGTTCAAT | gRNA for KD |
| SETX-shRNA-2 | AAGGATGATGCTCTTGCTAAAT | gRNA for KD |
| SETX-SP1 | CGTTCATGTAGAAGCAAGTANGG | gRNA for KO in hiPSCs |
| SETX-SP2 | TATTTCAGTATTTCAAGAAANGG | gRNA for KO in hiPSCs |
| SETX-SP3 | TGCCCAGTGATGTCAAACAGNGG | gRNA for KO in hiPSCs |
| SETX-SP4 | TGTTGAAGCACTTTGTCGGA | gRNA for KO in Fibroblasts |
| OCT3/4-F | CGAGAGGATTTTGAGGCTGC | RT-PCR |
| OCT3/4-R | CGAGGAGTACAGTGCAGTGA | RT-PCR |
| SOX2-F | AGGATAAGTACACGCTGCCC | RT-PCR |
| SOX2-R | TTCATGTGCGCGTAACTGTC | RT-PCR |
| NANOG-F | TGTCTTCTGCTGAGATGCCT | RT-PCR |
| NANOG-R | CAGAAGTGGGTTGTTTGCCT | RT-PCR |
| KLF4-F | TCTCCAATTCGCTGACCCAT | RT-PCR |
| KLF4-R | CGGATCGGATAGGTGAAGCT | RT-PCR |
| PAX6-F | GGGCGGAGTTATGATACCTACA | RT-PCR |
| PAX6-R | ATATCAGGTTCACTTCCGGGAA | RT-PCR |
| OTX1-F | TACGCCCTCCTCTTCCTACT | RT-PCR |
| OTX1-R | GCATGTGGGTGGTGATGATG | RT-PCR |
| DCN-F | CTGAAGAACCTTCACGCATTGA | RT-PCR |
| DCN-R | GGCAATTCCTTCAGCTGATTCT | RT-PCR |
| IGF2-F | CAATATGACACCTGGAAGCAGT | RT-PCR |
| IGF2-R | GTAGAGCAATCAGGGGACGG | RT-PCR |
| GATA2-F | ACCTGTTGTGCAAATTGTCAGA | RT-PCR |
| GATA2-R | ATCCCTTCCTTCTTCATGGTCA | RT-PCR |
| SOX7-F | ACTCCACTCCAACCTCCAAG | RT-PCR |
| SOX7-R | TTCATTGCGATCCATGTCCC | RT-PCR |
| SOX17-F | ATCGGGGACATGAAGGTGAA | RT-PCR |
| SOX17-R | TCCTTAGCCCACACCATGAA | RT-PCR |
| GAPDH-F | CAAATTCCATGGCACCGTCA | RT-PCR |
| GAPDH-R | GGACTCCACGACGTACTCAG | RT-PCR |

**Note: Tables S4-S9 were uploaded as Excel files separately.**

**Table S4. DEGS and GO_MN (KO vs WT).** Differentially expressed genes in WT and SETX KO iPSC-derived motor neurons (MNs) at 4 days post-viral infection, and Gene Ontology (GO) analysis results.

**Table S5. DEGs and GO_NPC (KO VS WT).** Differentially expressed genes in WT and SETX KO iPSC-derived neuronal progenitor cells (NPCS), and Gene Ontology (GO) analysis results.

**Table S6. DEGs and GO_WT (MN vs NPC).** Differentially expressed genes in WT MNs and NPCs, and Gene Ontology (GO) analysis results.

**Table S7. DEGs and GO_KO (MN vs NPC).** Differentially expressed genes in SETX KO MNs and NPCs, and Gene Ontology (GO) analysis results.

**Table S8. DEGs and GO_NPC-MN transition (KO vs WT).** Integration of analysis of DEGs during NPC-MN transition in WT and SETX KO, and Gene Ontology (GO) analysis results.

**Table S9. DEPs MN (KO vs WT).** Differentially expressed proteins (DEPs) in WT and SETX KO iPSC-derived motor neurons (MNs) at 4 days post-viral infection.
